# The circular RNA Ataxia-telangiectasia mutated (*cATM*) regulates oxidative stress in smooth muscle cells in expanding abdominal aortic aneurysms

**DOI:** 10.1101/2022.11.11.516115

**Authors:** F. Fasolo, G. Winski, Z. Li, Z. Wu, H. Winter, N. Glukha, J. Roy, R. Hultgren, J. Pauli, A. Busch, N. Sachs, C. Knappich, H.H. Eckstein, R.A. Boon, V. Paloschi, L. Maegdefessel

## Abstract

An abdominal aortic aneurysm (AAA) is a pathological widening of the aortic wall characterized by loss of smooth muscle cells (SMCs), extracellular matrix degradation, and local inflammation. This condition is often asymptomatic until rupture occurs, leading to high morbidity and mortality rates. Diagnosis is often accidental, and for now, the only available treatment option remains surgical intervention. Circular RNAs (circRNAs) are RNA loops that originated from backsplicing, which have received increasing attention as a novel class of functional non-coding RNAs contributing to cardiovascular physiology and disease. Their high structural stability, combined with a remarkable enrichment in body fluids, make circRNAs promising disease biomarkers. We aimed to investigate the contribution of circRNAs to AAA pathogenesis and their potential application as biomarkers for AAA diagnosis. Combined circRNA array and quantitative real-time PCR analysis revealed the presence of differentially expressed circular transcripts stemming from AAA-relevant gene *loci*. Among these, the circRNA to the Ataxia-telangiectasia mutated gene (c*ATM*) was upregulated in human AAA tissue specimens, in AAA patient-derived SMCs, and serum samples collected from aneurysm patients. In control primary aortic SMCs, c*ATM* increased upon angiotensin II stimulation, while its silencing triggered apoptosis. Furthermore, doxorubicin could induce cATM expression, supporting a link with acute stress response in SMCs. Constitutively higher c*ATM* expressing AAA patient-derived SMCs were less vulnerable to oxidative stress-induced cell death and survival pathways enriched when compared to control SMCs. Taken together, this data supports the role of *cATM* in adapting SMCs to oxidative stress in the vascular AAA micromilieu. This molecular signature provides an additional parameter to be included in procedures for AAA screening in combination with already established practices.

## INTRODUCTION

Abdominal aortic aneurysms (AAAs) are defined as localized pathological widening of the aortic diameter to more than 1.5 times its normal size^1^. Aortic aneurysms primarily affect the infrarenal aorta and are more prevalent among men over 65 years of age. Besides gender and age, risk factors include tobacco use and positive family history, suggesting a genetic association. On the molecular level, extensive loss of smooth muscle cells (SMCs) has been shown to be accompanied by metalloproteinases-mediated extracellular matrix degradation and local inflammation^2^. By further triggering metalloproteinase activity and SMC apoptosis, oxidative stress exacerbates the pathophysiology of inflammation, which may ultimately lead to thrombus formation^3^. Vessel dilation is often progressive, and AAAs remain mostly asymptomatic unless an acute rupture of the aortic wall occurs, accompanied by high mortality rates^4^. To date, established AAA prognostic indicators are still missing, making repeat imaging to monitor AAA expansion necessary. Furthermore, due to the lack of effective pharmacological treatments, the only available therapy is surgical intervention, either *via* an endovascular stenting approach (EVAR) or open repair (OR). The critical threshold for intervention is reached when the aortic diameter is found to be greater than 5.5 cm^5^. Identifying alternative, less invasive therapeutic strategies, relies on a deeper comprehension of the molecular mechanisms underlying AAA development and progression.

In the past decade, circRNAs have received increasing attention as a novel class of functional RNA molecules regulating biological processes’ physiological and pathological aspects. CircRNAs are covalently closed RNA loops generated from backsplicing events in which a downstream 5’ splice site (ss) is joined and ligated with an upstream 3’ ss^6^. They can be transcribed from either strand of exons (exonic circRNAs - the most abundant class) or introns (intronic circRNAs) of host genes. Circularization is often triggered by the presence of repetitive elements in backsplicing junction-flanking introns^7^. Although poorly expressed, their class-specific structural features confer higher stability than messenger RNAs (mRNAs), as the lack of a free –OH end makes them highly resistant to RNAseR degradation^8,9^. According to their subcellular localization, circRNAs contribute to the regulation of gene expression through multiple mechanisms. They participate in splicing, may act as miRNA or protein ‘sponges’, and can interfere with the pre-mRNA processing machinery^10^. CircRNAs can be secreted in body fluids, including saliva and serum, making them ideal candidate biomarkers in clinical practice^11^.

In the context of cardiovascular disease (CVD), circular *ANRIL* (circ*ANRIL*) was the first circRNA to be associated to the molecular regulatory mechanisms at the basis of CVD risk phenotype at the cyclin-dependent kinase inhibitor 2A/B (*CDKN2A/B*) *locus*^12,13^. As of now, many circRNAs have been shown to significantly contribute to CVD pathophysiology ^14,15^. The SMC-enriched circRNA circChordc1, for example, protects from AAA pathology by promoting SMCs contractility and improving their survival^16^. cZNF292 contributes to the maintenance of the structural integrity of the aortic endothelium by regulating flow responses in endothelial cells^17^.

For this current study, we profiled differentially expressed circRNAs in AAAs *vs*. non-AAA human tissue specimens. Next, we investigated the role of the most promising candidates in AAA pathogenesis and progression and finally explored their potential application as a diagnostic AAA biomarker.

## RESULTS

### Circular RNA profiling in human abdominal aortic aneurysm (AAA) disease

To evaluate the role of circRNAs in AAA pathogenesis and disease progression, we performed a microarray analysis on a cohort of 17 human tissue specimens, including 6 non-AAA controls (CTRL, obtained from organ transplant surgery) and 11 AAA aortas collected from elective open repair (OR) surgery. The employed array chips (Arraystar, #AS-S-CR-H-V2.0) display all known annotated human circRNAs (13.617 in total, **Figure S1B**), with array probes targeting circRNA-specific junctions. Interestingly, we discovered that a considerable amount of circRNA transcripts were significantly up-or downregulated in diseased tissue (**Figure 1A; Figure S1A**). Most deregulated targets in our dataset originated from circularization of one or multiple mRNA exons (**Figure 1B**). Interestingly, among circRNA-cognate mRNAs, gene ontologies (GO), like regulation of transcription and SMCs proliferation, caught our attention, as they are established mechanistic drivers in AAA development (**Figure 1C**). As exonic circRNAs have previously been shown to contribute to regulating host gene expression^18^, the relevance of the linear counterpart (mRNA) in AAA pathophysiology was used as a criterion to shortlist circRNA candidates for downstream validation experiments.

**Figure 1.**
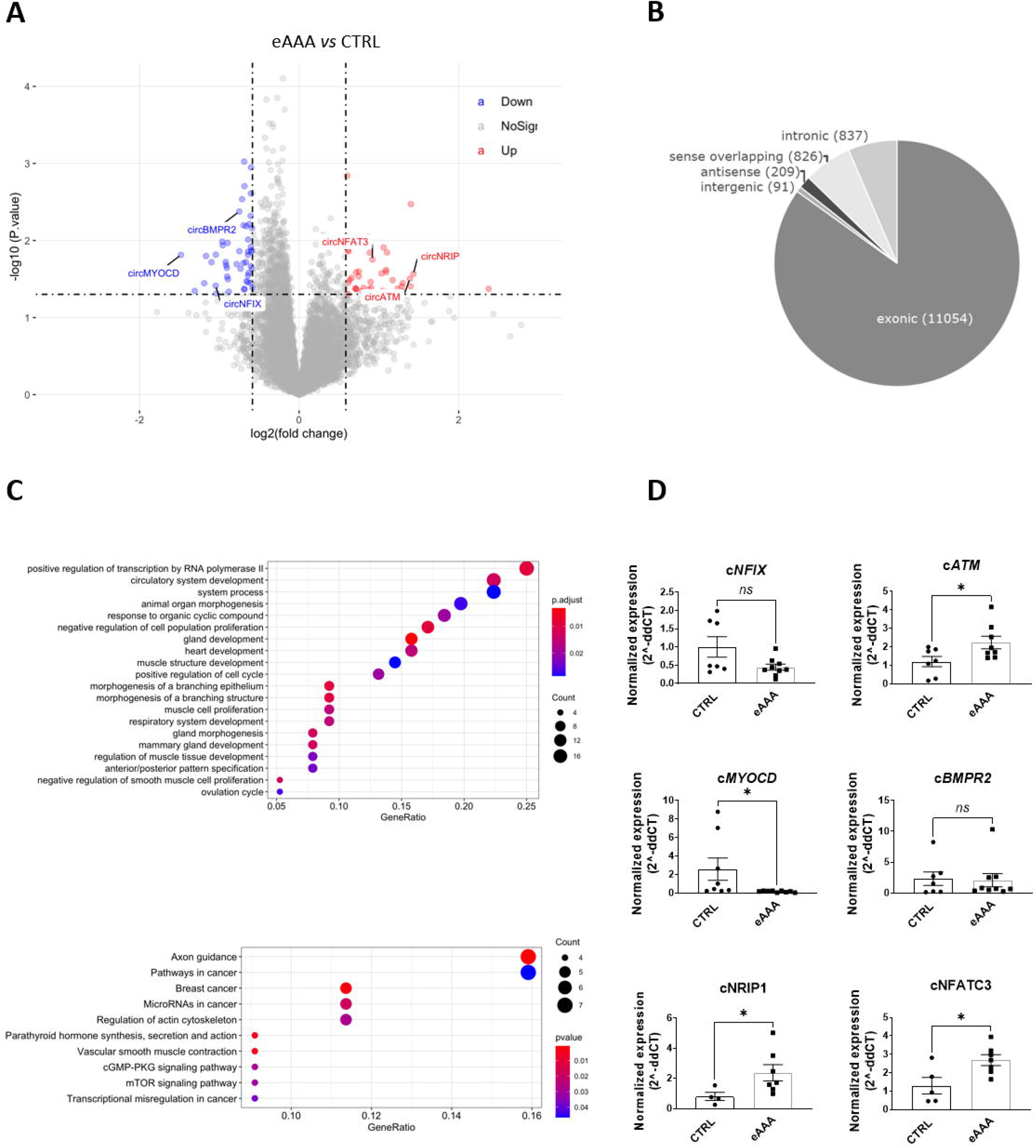
Circular RNAs (circRNAs) are de-regulated in human abdominal aortic aneurysm (AAA). **A**. Volcano plot depicting down-(left, blue) and up-(right, red) regulated circRNAs in human elective AAA (eAAA) *versus* control (CTRL) aorta specimens, as resulted by array experiments. Log2 fold change and -log10 p-value are plotted on the x and y-axis, respectively. IDs of validated circRNAs are highlighted. **B**. Pie chart illustrating the proportion of exonic (89.8%), intronic (5.7%), sense-overlapping (3.4%), and antisense (1.1%) array-identified differentially expressed circRNAs. Absolute numbers are further indicated for each group. **C**. Top: Gene Ontology (GO) enrichment analysis of linear mRNA counterparts of differentially expressed exonic circRNAs. Only significantly enriched (adjusted-p < 0.05) GO-terms are shown. Bottom: Heat-map of circRNA counterpart expression within the enriched GO-terms. **D**. Quantitative real-time PCR (qRT-PCR) validation of c*ATM* (hsa_circ_0003641), c*NFIX* (hsa_circ_0005660), c*BMPR2* (hsa_circ_0003218), c*MYOCD* (hsa_circ_0042103), c*NRIP1* (hsa_circ_0005660) and c*NFATC3* (hsa_circ_0004771) differential expression in human AAA and CTRL aortas. 2^(-ddCT) was calculated by normalizing on RPLPO. Data are represented as mean ± SEM. Statistics: T-test. p-values < 0.05 was considered significant. Abbr.: NS= non significant; eAAA= elective.

To prove circularity, the selected targets were amplified with divergent primers in RNAseR-treated samples, and the specificity of the resulting amplicons was checked by Sanger sequencing. 6/12 backsplicing junctions were validated and submitted to downstream analysis (**Figure S2A**). Resistance to RNAseR digestion was assessed *via* comparison with a linear mRNA (*ACTA*2) in both AAA tissue (**Figure S2B**) and human aortic SMCs (hAoSMCs) (**Figure S2C**). Furthermore, subcellular localization of both circular and linear variants was profiled in hAoSMCs (**Figure S2D**). Globally, circRNAs were predominantly detected in the cytosolic compartment, even though considerable amounts were also present in the nucleus (except for *NRIP*, for which this trend was reverted). QRT-PCR validation was eventually performed for the following targets: hsa_circ hsa_circ_0005660 (c*NFIX*), hsa_circ_0003641 (c*ATM*), hsa_circ0042103 (c*MYOCD*), hsa_circ003218 (c*BMPR2*), hsa_circ0005615 (c*NRIP1*) and hsa_circ0004771 (c*NFATC3*). Array results were confirmed for all circRNA targets, except for circ*BMPR2*, for which no significant difference could be observed (**Figure 1D**). Of notice, most of the host genes of validated circular targets (**Figure S3**) encoded for transcription factors (kinases), and interestingly corresponding deregulation trends mirrored the ones detected for circRNA (except for *NFIX*, **Figure S4**). In summary, differential expression of circRNAs in AAA *vs*. CTRL human aortic tissue specimens was accompanied by similar deregulation of the cognate mRNAs.

### c*ATM* is increased in human AAAs, and its inhibition triggers *ATM* mRNA expression and apoptosis in smooth muscle cells

Inflammatory mediators and radical oxygen species (ROS) released by infiltrating immune cells contribute to oxidative stress by inducing DNA damage, which triggers apoptotic cell death within the AAA micromilieu. *ATM* is a ubiquitously expressed nuclear protein sensing double-strand break (DSB) and coordinating the DNA damage response (DDR). According to the extent of the damage, survival or apoptotic gene programs are induced. Activated by autophosphorylation, ATM phosphorylates downstream p53, responsible for the initiation of the apoptotic cascade. We hypothesized that aberrant *ATM* expression could affect AoSMCs resistance to oxidative stress and be thus linked to their loss. Immunohistochemistry (IHC) on human aneurysmal sections revealed ATM signals in the *adventitia* and in the *media*, where SMCs reside (**Figure S5A**). We first verified ATM and smooth muscle cell L-actin (LSMA) co-expression on human AAA tissue samples (**Figure 2A, Figure S5B**). Single-cell RNA sequencing (scRNA-seq) analysis carried out from 4 human AAA specimens confirmed ATM expression in vascular mural structural cells (**Figure 2B, C; Figure S6**). On the other hand, combined *in situ* hybridization (ISH) and LSMA staining on consecutive AAA sections indicated the presence of *cATM* in SMCs (**Figure 2D and Figure S5C, D**), further corroborated by ISH in primary AoSMCs (**Figure S5E**). The circRNA was predominantly localized in the cytoplasm, even though a considerable amount was also present in the nuclear compartment (**Figure 2D, Figure S2D**).

**Figure 2.**
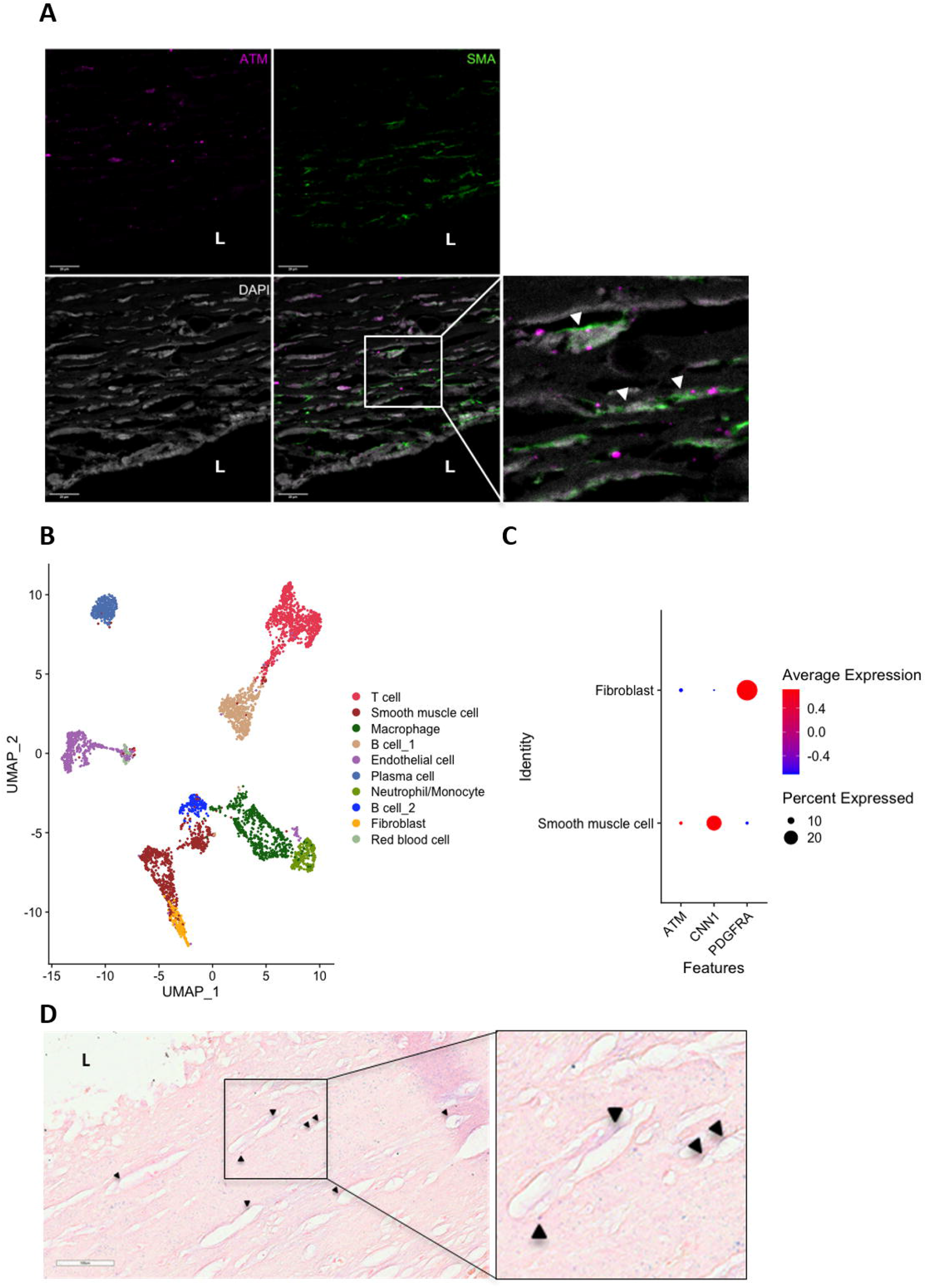
*ATM* and circular *ATM* (*TM*) are expressed in human AAA smooth muscle cells (AoSMCs). **A**. Immunofluorescence on human AAA section showing co-localization of ATM (purple) and LSMA (green) signal. Imaging was carried out with confocal microscopy. The zoomed area includes double-positive cells (white arrows). **B**. Uniform Manifold Approximation and Projection (UMAP) plot shows the cell clusters identified by single-cell RNA sequencing (scRNA-seq) of n=4 human AAA tissue specimens. Dot red line highlights mural structural cell clusters. **C**. Dot plot showing relative ATM expression in mural structural cell clusters (Fibroblasts) and SMCs) with respective expression profiles of one representative cluster’s marker (CNN1 and MYOCD). Dot color represents the average expression level (blue=low, red= high) and dot size depicts the percentage of cells expressing the gene in a given cluster. **D**. *In situ* hybridization (ISH) of c*ATM* (purple signal, black arrows). ISH was performed with miRCURY LNA miRNA ISH Optimization Kits for FFPE sections; cATM probe was designed on the backsplicing junction. Abbr.: L= lumen

Next, we focused on c*ATM*-mediated regulation at the *ATM locus* and profiled *ATM* and *cATM* expression patterns in disease. In patients’ AAA specimens, c*ATM* and *ATM* mRNA upregulation (**Figure 1D; Figure S7A**) was accompanied by reduced global ATM protein levels (**Figure S7B**). In line with this, *in vitro* AAA-mimicking angiotensin II (AngII) stimulation caused a remarkable c*ATM* increase (**Figure 3A**). We then took advantage of primary AoSMCs isolated from AAA biopsies. Compared to commercially available control (CTRL) AoSMCs provided by a healthy donor, patient-derived cells displayed higher c*ATM*, but lower *ATM* mRNA levels (**Figure 3B**). To better elucidate the impact of c*ATM* in regulating host gene expression in AoSMCs, we conducted silencing experiments. We monitored both circular and linear *ATM* in CTRL and patient-derived primary cells. We designed two alternative siRNAs targeting c*ATM* (**Figure S8A**) and observed that only sic*ATM*2 (and not sic*ATM*1) selectively targeted circular (**Figure S8B, left**) and not *ATM* mRNA (**Figure S8B, right**). Upon sic*ATM*2, in both AAA-derived and CTRL AoSMCs, *ATM* mRNA was significantly increased after 72h (**Figure 3C, 3D**), although this did not affect ATM protein levels (**Figure S8C, D**). In CTRL AoSMCs, silencing resulted in increased apoptosis, as measured by Caspase 3/7 live imaging (**Figure 3E**) and Caspase 3/ Active Caspase 3 Western Blotting (WB) signal (**Figure 3F**). In addition, decreased expression of the survival gene *BCL2* was observed upon c*ATM* Knock-down (KD) (**Figure 3F**). In summary, increased *cATM* expression detected in AAA tissue specimens was confirmed in AoSMCs, *via* comparing CTRL *versus* patient-derived cells. On the contrary, opposing trends were observed for *ATM* mRNA, upregulated in AAA tissue, but downregulated in patient-derived AoSMCs. Moreover, in CTRL AoSMCS, c*ATM* depletion induced apoptosis while down-regulating pro-survival genes.

**Figure 3.**
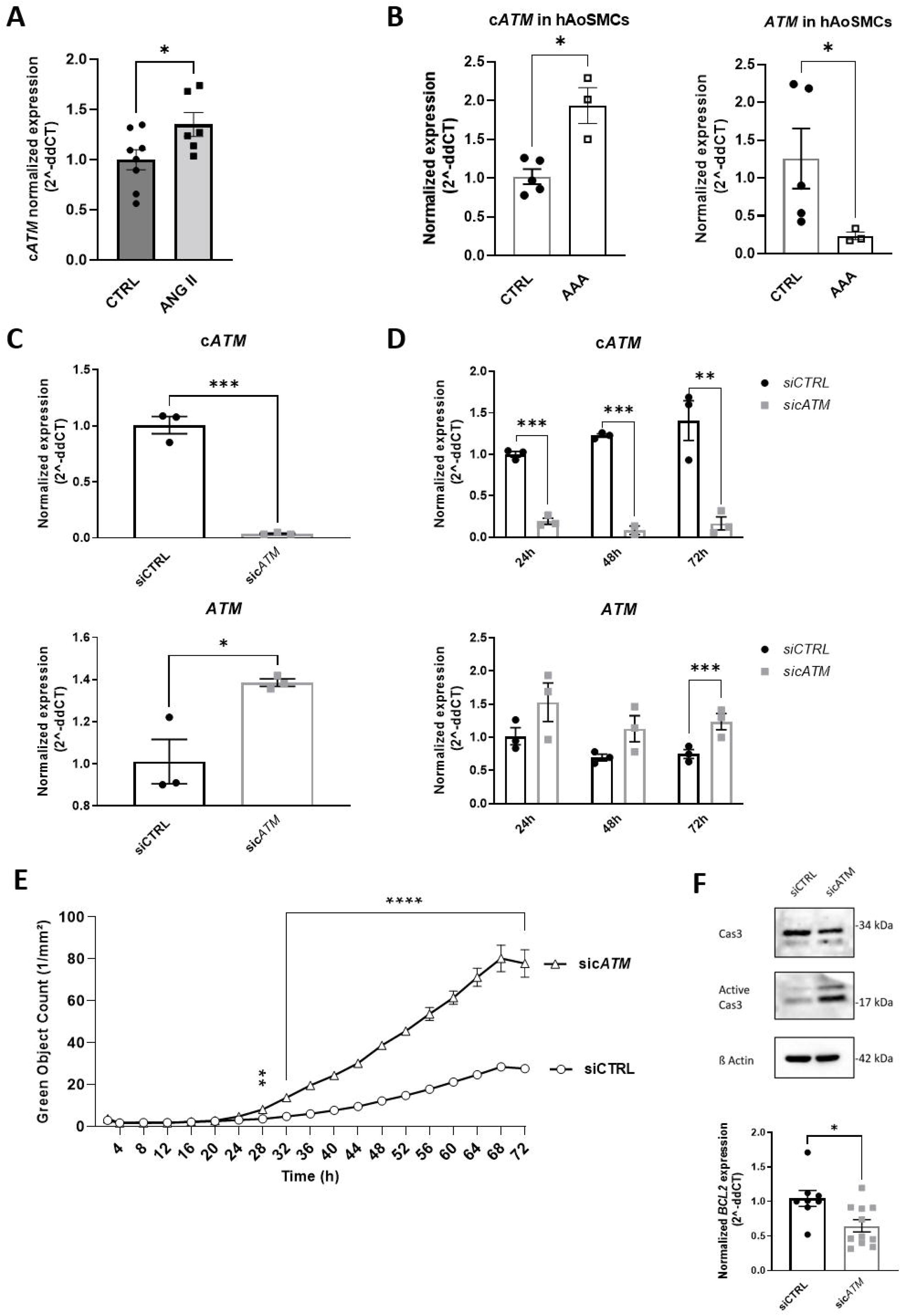
Modulation of gene expression at *ATM locus* in AoSMCs. **A**. Control (CTRL) hAoSMCs to treatment with 0.2μM Angiotensin II for 24h and *cATM* expression measured by qRT-PCR. Water served as control treatment **B**. Comparison of c*ATM* (left) and *ATM* mRNA (right) expression in control donor (CTRL) *versus* AAA patient-derived hAoSMCs (AAA) by qRT-PCR. AAA (**C**) and CTRL hAoSMCs (**D**) were transfected with 100nM sic*ATM*2 and c*ATM* and *ATM* mRNA expression measured by qRT-PCR at different time points. 2^(-ddCT) was calculated by normalizing on RPLPO. KD effects on apoptosis were measured by live imaging and quantification of fluorescent Cas3/7 (**E**) and WB (**F**) on Cas3/Active Cas3 protein. *BCL2* expression was further monitored as a marker of cell survival (**F**). N=3. Data are represented as mean ± SEM. Statistics: T-test. p-values < 0.05 was considered significant. Abbr.: Ang II= angiotensin II.

### Higher c*ATM*-expressing AAA patient-derived AoSMCs are more resistant to oxidative stress

Once demonstrated that modulation of c*ATM* expression affected *ATM* mRNA levels and CTRL AoSMCs survival, we hypothesized that c*ATM* upregulation in patient-derived cells could be part of an early stress response triggered by the AAA environment. As oxidative stress represents a major hallmark of the AAA micromilieu, we recreated a oxidative stress-rich climate, by treating CTRL and AAA patient-derived AoSMCs with doxorubicin or DMSO, as negative control. While c*ATM* levels in AAA-derived AoSMCs remained unaltered (**Figure 4A, up-left**), CTRL cells showed an augmented expression of this circRNA (**Figure 4A, up-right**) similar to what we observed in AAA tissue specimens. *ATM* mRNA was downregulated (**Figure 4A, down**) and induced by doxorubicin. *ATM* and p53 phosphorylation were monitored as treatment control (**Figure 4A, right**). Noteworthy, combined live apoptosis assay and WB pointed out lower apoptotic rate upon stress in patient-derived AoSMCs compared to CTRL AoSMCs (**Figure 4B**). Although likely to be triggered by stress, higher c*ATM* expression characterizing patient-derived cells seemed beneficial rather than detrimental. To investigate this hypothesis, we sequenced RNA extracted from both primary cells. As expected, a SMC typical gene signature was detected in both AAA-derived and CTRL SMCs (**Figure S9**). Of notice, expression of *MYOCD*, a transcription factor contributing to determining the contractile SMC phenotype, was higher in CTRL cells. No substantial differences were detected when assessing the expression of synthetic markers, such as COL5A2, FBN1, VIM, and FN1 **(Figure S9)**. According to Reactome and Gene Ontology (GO) analysis, proliferation-related pathways and cell cycle-associated ontologies appeared greatly enriched in AAA patient-derived cells compared to CTRL (**Figure 5A, Table S1**). To our surprise, the main difference between the two primary cell lines could be detected in cell cycle/proliferation-related gene expression patterns, highlighting a more vital proliferative state in AAA-derived AoSMCs (**Figure 5B**). In addition, AAA patient-derived AoSMCs were further characterized by the enrichment of ATM-linked cell functions and signaling, including replication stress, DNA repair and p53-transduction (**Figure 5A**). To further assess this in more detail, we subsequently monitored differences in expression profiles of genes involved in the ATM pathway and apoptosis (**Figure 5B**). First, decreased *ATM* levels in AAA *vs*. CTRL cells could be confirmed. Noteworthy, expression of *ATM* downstream effectors leading to activation of cell cycle checkpoints (*RAD9A)*, cell cycle arrest (*CDK2*), and DNA repair (*BRCA1*) pathways were predominantly detected in patient-derived cells. Moreover, survival pathways (*BCL2*) were enriched over apoptotic ones (*BAX*). Conversely, in CTRL AoSMCs, the expression of late apoptosis stages-related genes (*CAS3, BID*) was more pronounced. Furthermore, WB revealed that CTRL cells had higher default levels of phosphorylated ATM and p53 (**Figure 5C**). Taken together, these data suggest that higher c*ATM*-expressing AAA patient-derived AoSMCs are characterized by a greater resistance to oxidative stress, a more efficient DDR, and diminished apoptosis.

**Figure 4.**
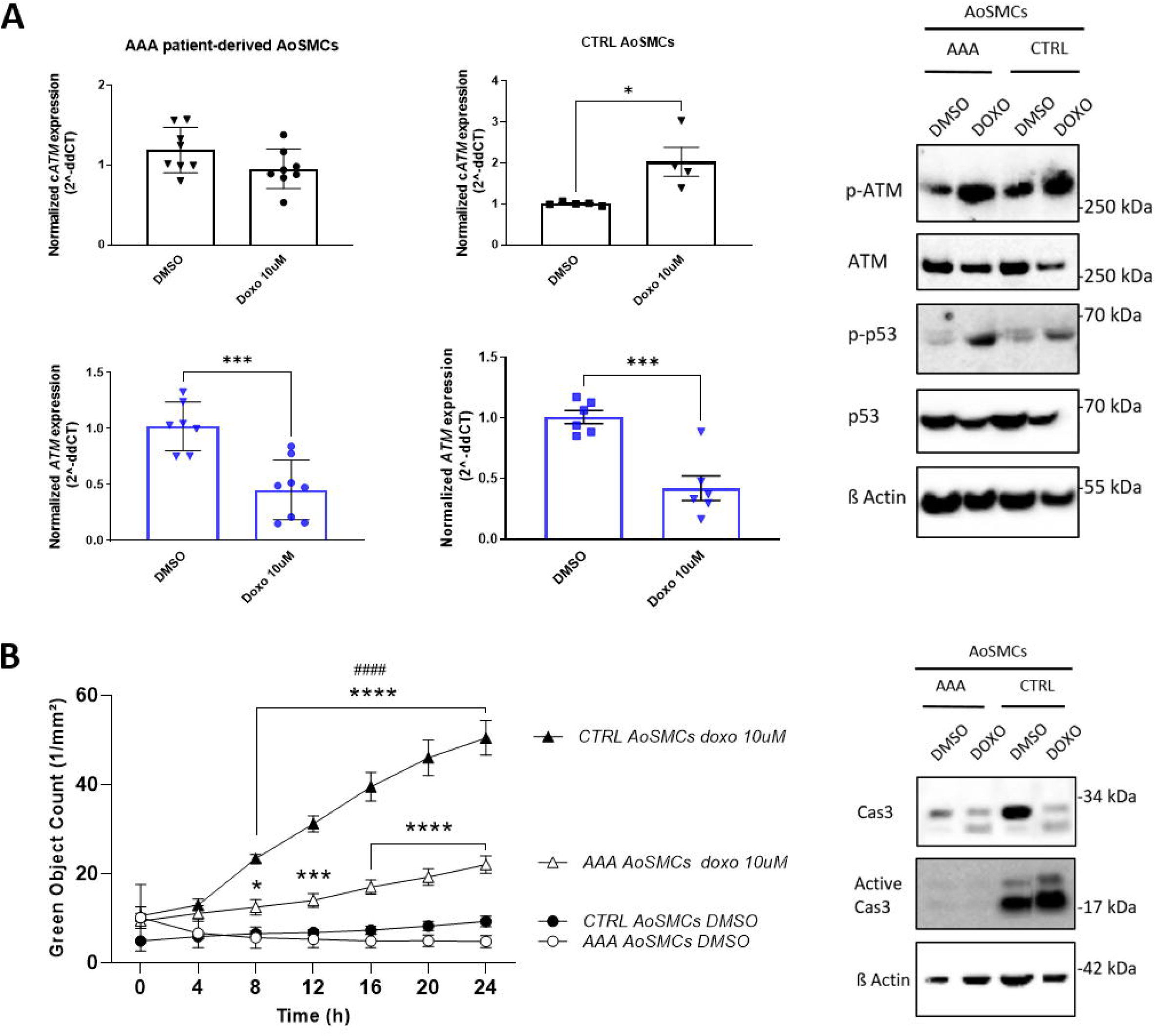
Effects of doxorubicin-induced oxidative stress in AAA patient-derived *vs* CTRL AoSMCs. **A**. AAA patient (left) and control donor-derived AoSMCs (right) were treated with 10 μM doxorubicin/ DMSO for 24h and c*ATM* (black) and *ATM* mRNA (blue) expression measured by qRT-PCR (left). 2^(-ddCT) was calculated by normalizing on *RPLPO*. ATM, phospho-ATM, p53 and phospho-p53 protein levels were also compared by WB (right). **B**. Treatment effects on apoptosis were measured by live quantification of fluorescent Cas3/7 and WB on Cas3/ Active Cas3 protein. (*) refers to comparison between treated/untreated CTRL or AAA-patient derived SMCs; (#) refers to comparison between treated CTRL vs AAA patient derived SMC. N=3. Data are represented as mean ± SEM. Statistics: T-test. p-values < 0.05 was considered significant. Abbr.: Doxo= doxorubicin.

**Figure 5.**
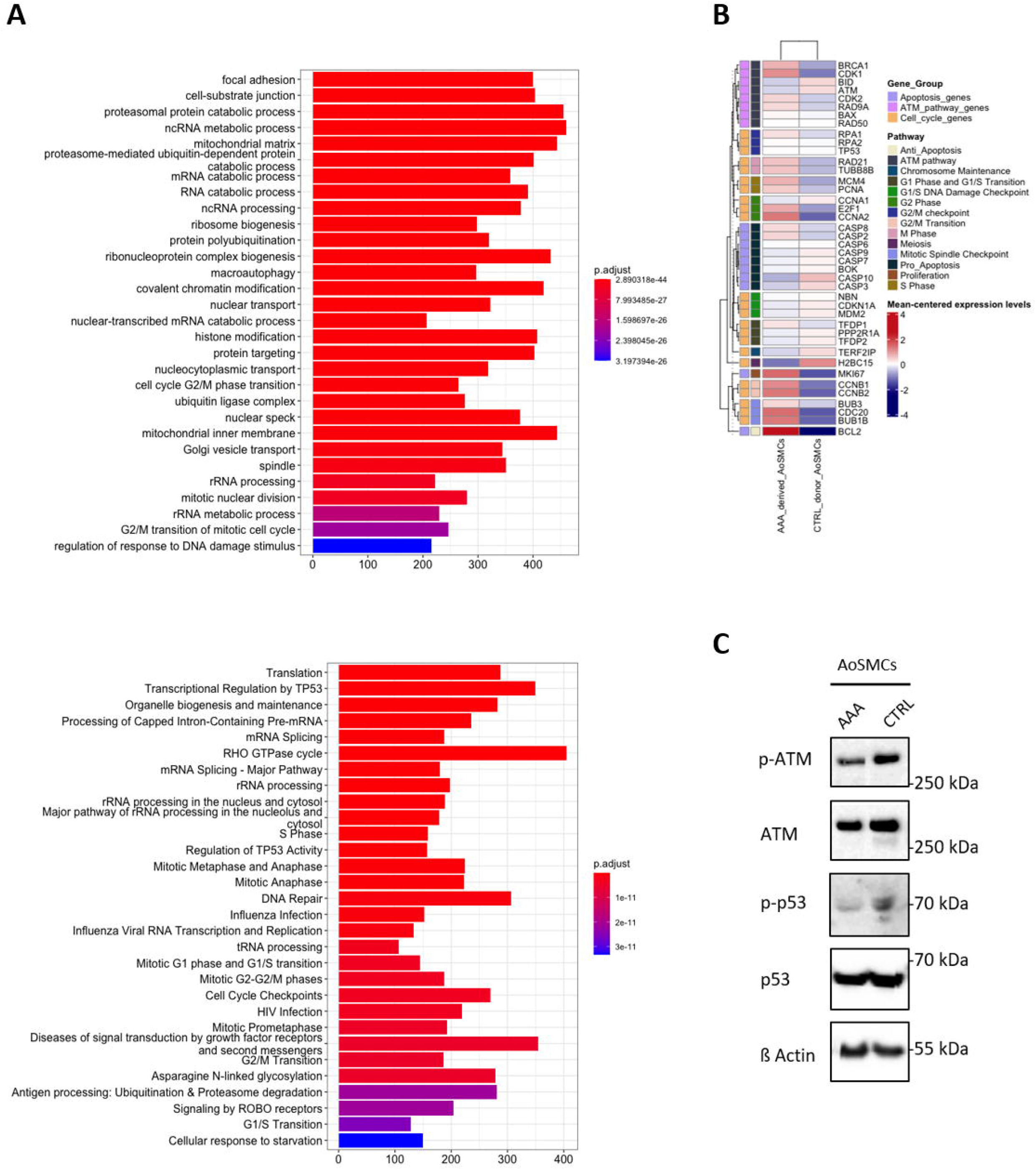
Expression profile of *ATM* pathway, apoptosis and cell cycle-related genes in AAA patient-derived *vs* CTRL AoSMCs. **A**. Top: Reactome (top, https://reactome.org/)- or Gene Ontology (GO, bottom, https://geneontology.org/) based gene set enrichment analysis (GSEA) was performed to determine whether sets of genes involved in pathways/GO-terms of interest tend to be over-represented at the extremes of an ordered list of genes. This was created by ranking all genes according to a statistic, from high to low. The statistical significance (nominal p-value) of the over-representation was evaluated using a method based on an adaptive multi-level split Monte Carlo scheme. The top 30 significant pathways/GO-terms with qvalue (FDR-adjusted p-value) < 0.05 were included. **B**. Heatmap depicting expression profiles of apoptosis-, ATM pathway- and proliferation-related genes in AAA patient (left) and control donor-derived AoSMCs (right). **C**. Western immunoblot comparing phosphorylation status of ATM pathway proteins (ATM and p53) in AAA patient-(left) and control donor-derived AoSMCs (right).

### c*ATM* can be detected in serum of AAA patients and elevated with disease

A crucial aspect for the application of circRNA in clinical practice is their suitability to be employed as biomarkers due to their presence and structural resistance to degradation in body fluids (*e*.*g*., blood and saliva). By taking advantage of serum samples collected from 9 AAA patients undergoing elective surgery (eAAA) and 10 age/sex-matched controls (CTRL) with peripheral artery disease (PAD), we compared c*ATM* expression in both groups. Circulating c*ATM* was significantly higher in AAA patients (**Figure 6A**), supporting that a certain range of c*ATM* expression might reflect the disease status. Next, we aimed at detecting ATM protein in blood. Interestingly, ATM levels were not significantly different between AAA and control (PAD) patients (**Figure 6B**), suggesting that only the circular isoform is suited to diagnose patients with aortic aneurysms.

**Figure 6.**
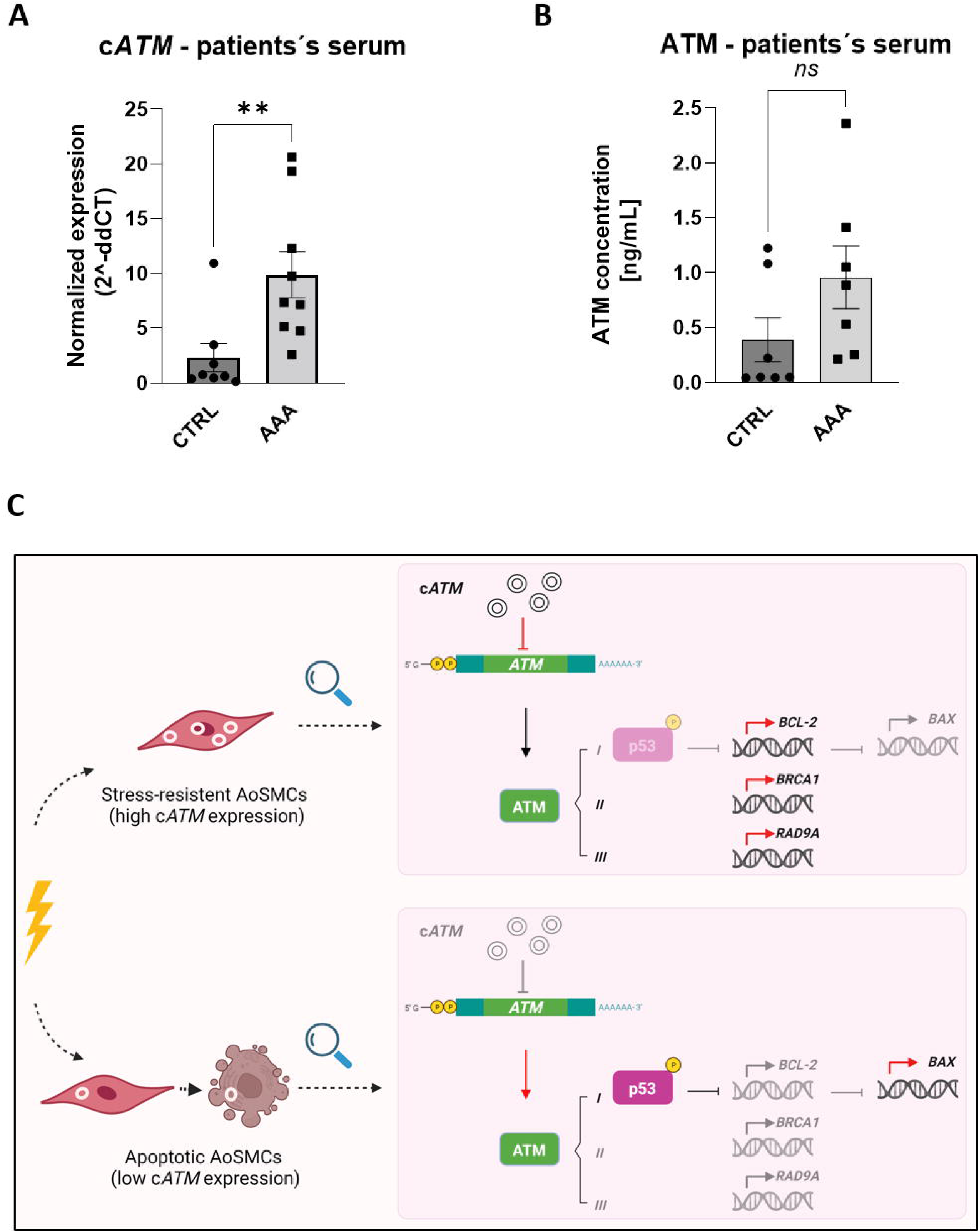
Circulating c*ATM* as AAA biomarker. **A**. RNA was extracted from serum samples collected from AAA patients and peripheral artery disease (PAD) controls and submitted to qRT-PCR to monitor c*ATM* expression. 100ng of an *in vitro* transcribed *GFP* RNA were spiked in to every sample and used as a reference to calculate 2^(-ddCT). **B** Sandwich ELISA was performed to monitor the expression of ATM protein in patients’ serum. ATM concentration was calculated based on the standard curve. In both charts, data are represented as mean ± SEM. Statistics: Mann-Whitney U. p-values < 0.05 was considered significant. **C**. Proposed mechanism of c*ATM*-mediated regulation of *ATM* expression and its effects on survival/ apoptosis in human AoSMC.

## DISCUSSION

Abdominal aortic aneurysm is an asymptomatic condition accompanied by a high mortality rate related to its acute rupture risk. In Western countries, the annual incidence of new AAA diagnoses, which despite screening programs in some countries, are mostly accidental, is approximately 0.4 to 0.67 percent^19-22^. A clear disease etiology has yet to be established. However, the mechanisms at the basis of the development, dilation, and rupture of AAA have long been the object of scientific investigations. The anatomical characteristics and diameter of AAAs remain the only parameters to estimate individual risk of rupture and the timing of surgical intervention. As the natural history of AAA varies among patients, an accurate diagnosis would strongly benefit from specific molecular predictors, which in combination with other known risk factors (*e*.*g*., age, male sex, smoking status, blood pressure, and dyslipidemia) could improve diagnosis and patient management.

In search for unexplored AAA gene expression signatures, we focused our attention on circRNAs. These are covalently closed RNA loops already described in the late 1970s in plants viroids^23^. Only more recently it was proven that they can exert critical regulatory functions in development and disease^24,25^. In addition to the regulation of gene expression at multiple levels and through multiple mechanisms, circRNAs are tissue-specific and can be secreted into blood and saliva. Their circular structure, ensuring stability, makes them particularly suitable as biomarkers^26^. One criterion a candidate diagnostic biomarker has to meet is biological plausibility, establishing a cause-and-effect relationship between a natural factor and a particular disease^27^. Taking all of this into consideration, our study aimed at *i)* profiling circRNAs expression in AAA *vs*. control human specimens, *ii)* selecting disease-relevant targets associated with functional phenotype, and *iii)* addressing their clinical application as biomarkers.

To detect AAA-characterizing circRNAs, we took advantage of microarrays with backsplicing junction-mapping probes specific for all annotated circRNAs. Enzymatic (RNAseR) linear RNA removal was applied prior to hybridization to enrich, capture, and quantify circular RNA targets with high sensitivity and specificity. Array-based quantification has the advantage of being less affected by the number of transcripts in individual samples compared to RNA sequencing. However, a relevant drawback of RNAseR treatment is the potential loss of an undefined portion of circRNAs species, which may undergo degradation more efficiently ^28^.

CircRNAs can be transcribed from different genomic regions and are accordingly distinguished into intergenic, intronic or exonic. Exonic circRNAs share their exons with mRNAs and constitute the prevalent subgroup. In line with this, most differentially expressed circRNAs in our dataset had a linear mRNA counterpart. A growing body of evidence supports that non-coding circular transcripts affect gene expression at the *locus* of origin. Regulation may be achieved through the process of circularization itself^29^, or alternatively, circRNAs can be provided with specific functional features (reviewed in^30^). We prioritized validation of circRNA targets having a linear mRNA counterpart, whose deregulation affected cell proliferation, stress response, and apoptosis, which have a central role in AAA pathogenesis. mRNA expression of ataxia telangiectasia (*ATM*, a key regulator of the p53-mediated DNA damage response) is increased in tricuspid aortic valve (TAV) and in bicuspid aortic valve (BAV)-associated thoracic aortic aneurysms (TAA) specimens and accompanied by a pro-proliferative state^31^. Different Malan syndrome-characterizing nuclear factor I X (*NFIX*) mutations have been previously associated with early-onset thoracic aortic aneurysm development^32^. Myocardin (*MYOCD*) has been extensively acknowledged as an essential transcription factor for maintenance of the SMCs contractility^33^. The bone morphogenetic protein (*BMP*) signaling is known to be tightly involved in pulmonary artery (PA) dilatation following pulmonary arterial hypertension (PAH)^34^. Macrophage nuclear factor of activated T-cell C3 (*NFATC3*) limits inflammation and foam cell formation^35^. The nuclear receptor interacting protein 1 (*NRIP1*) is a transcription factor modulating the transcriptional activity of the estrogen receptor.

Validation of array data in aortic tissue specimens confirmed differential expression of *cNFIX, cATM, cMYOCD, cNRIP1*, and *cNFATC3*. In line with most literature findings^36^, identified deregulation trends of circRNAs and respective linear counterparts were concordant. This would support the “microRNA sponge” functional model, where the upregulation of a given circRNA generates in turn, an increase of its cognate mRNA by sequestering and preventing specific microRNAs from silencing their targets. This could represent a mechanism to promptly provide the cell with plenty of ready-to-translate mRNAs to sustain the synthesis of essential proteins. Protein degradation is extensively observed in AAA, because of metalloproteases activity, immune cell infiltration, and inflammation-induced oxidative stress. ATM, one of the main PI3 kinases involved in sensing DSB and regulating the DDR, was strongly depleted in AAA tissue specimens. Conversely, c*ATM* and *ATM* mRNA were upregulated. We speculated that c*ATM* transcription could be induced by ATM protein down-regulation and that sponging of *ATM*-targeting miRNAs by c*ATM* would augment *ATM* mRNA available for translation. Supporting this hypothesis, c*ATM* contains binding sites for *ATM*-targeting miR-567, miR-4677 5p and miR-4778 5p. Given the importance of efficient regulation of the DDR in an oxidative stress-rich environment as found in AAA, we further dissected the effects of c*ATM*-mediated regulation of *ATM* expression. *ATM* is a multi-exonic mRNA (canonical form: 64 exons; 3,056 residues; 350KDa) transcribed from the *ATM locus* at chr11, presenting about 30 annotated coding splicing variants^37^. *Via* modulation of gene expression networks controlling cell fate, *ATM* eventually guides the cell towards survival or programmed cell death. In the case of induced apoptosis, ATM, activated by autophosphorylation, can in turn phosphorylate downstream p53, responsible for the initiation of the apoptotic cascade. ATM expression is sustained in immune cells, ensuring an efficient coordination of the DDR. These are indeed particularly exposed to continuous genomic insults, as a consequence of abundant ROS production^38^. We speculate that trends of *cATM/ATM* expression detected in the analysis of bulk tissue can be mostly due to the presence of immune infiltrates. At aneurysmal sites, indeed, immune cells are massively recruited and contribute to generate a stress-rich environment. As oxidative stress represents a major threat to the integrity of AoSMCs during AAA progression, we aimed at studying *ATM* gene expression regulation in this cell population, with a special focus on the role of the *ATM* circular isoform. We found that c*ATM* was upregulated in angiotensin II-treated CTRL AoSMCs (derived from non-AAA patients), and its expression levels in AAA patient-derived AoSMCs were significantly higher compared to non-dilated CTRL AoSMCs. Conversely to what was observed in bulk tissue, linear *ATM* expression was depleted in AAA cells compared to CTRL ones. In line with this, c*ATM* silencing resulted in higher levels of *ATM* in both CTRL and AAA patient-derived AoSMCs. This implies that the grammar underlying *cATM* regulation in AoSMCs differs from the “miRNA sponge” model. Interestingly, c*ATM* deprivation in CTRL AoSMCs was associated with higher apoptosis. Even if disease-related, higher expression levels of c*ATM* in AAA seemed to thus be protective, rather than detrimental. This was further corroborated by a higher tolerance to doxorubicin-induced oxidative stress in AAA compared to CTRL AoSMCs, in which apoptosis was considerably higher. Coherent with our findings, in their study on DDR in an inducible human neural stem cell line (ihNSCs), Carlessi *et al*. showed that ATM protein depletion attenuates the ionizing radiation-induced DDR, as shown by reduced phosphorylation of the ATM target p53 and, in turn, the p53-dependent apoptotic response^39^. Here, we propose that a negative c*ATM*/*ATM* regulatory loop regulates gene expression at the *ATM locus* in AoSMCs, similar to previous studies that reported a modulating role for circRNAs in apoptosis^40^.

It is widely acknowledged that inflammatory mediators contribute to phenotypic and functional alterations in AoSMCs during AAA development (reviewed in^41^). Stress-triggered c*ATM* transcription would thus work as a signal for activating survival pathways (like *BCL2*) by fine-tuning *ATM* expression levels. Although the molecular dynamics at the basis of c*ATM* transcription and the specific mechanism through which this circRNA inhibits the expression of its linear counterpart remain to be elucidated, our work shows that circRNA expression is crucial in determining and maintaining phenotypic variability among SMCs (**Figure 6C**). The different AoSMCs utilized in our study (AAA patient-derived *vs*. non-dilated organ donor controls; CTRL) expressed different levels of c*ATM*, and, accordingly, presented different phenotypic states of the diseased and non-diseased aorta. Although sequencing was performed on CTRL and AAA-derived cells from two individuals, we found substantial differences in the expression of stress response and apoptosis-related genes. Together with our functional experiments, this profiling further corroborated our hypothesis of having selected a population of “AAA-adapted” cells, with augmented resistance to stress.

We eventually assessed that *cATM* expression profiles could be exploited as a diagnostic tool accessible in circulating blood. Coherent with our previous findings, c*ATM* was indeed upregulated in serum samples collected from AAA patients. It would be challenging to claim that a circRNA could be *per se* used as a biomarker in AAA screening, but we are confident in stating that given their high stability and relatively low detection costs, circRNA could play an essential role in molecular AAA diagnostics in the future. Its suitability in determining high-risk AAA patients with an elevated likeliness of rupture would be very desirable. It would further be a valuable compliment to the current diagnostic gold standard of B-mode ultrasound imaging, which cannot predict the rapidness of AAA growth, and thus is only of value in a sole diagnostic setting.

## CONCLUSIONS

In summary, our work emphasizes the importance of investigating non-coding RNA-mediated gene expression regulation as a tool to improve our comprehension of AAA molecular dynamics. In particular, we are able to provide an example of a circRNA-disease-signature and propose c*ATM* transcription as part of an early stress response triggered by AAA-characteristic microenvironmental changes. By regulating gene expression at the *ATM locus* and contributing to the activation of survival pathways, c*ATM* upregulation functions as an ‘early warning signal’ that promotes a switch towards stress-resistant SMC phenotypes.

## METHODS

### Human samples collection and storage

Human samples employed in this study were provided by the Munich Vascular Biobank (MVB) and the Stockholm Aortic Aneurysm Biobank (STAAB). AAA tissue specimens or blood were collected from patients who underwent elective surgical repair in both hospitals. Aortic fragments harvested in organ explant procedures or serum from patients with peripheral artery disease (PAD) were used as respective controls. Tissue specimens were put in RNAlater for 24h, and then snap-frozen and stored at −80°C. All patients provided their written informed consent and in accordance with the Declaration of Helsinki. The studies were approved by the local Ethics committees. All patient samples were matched for age (average: 65,26 years) and sex (90% males).

### Circular RNA array experimental pipeline and analysis

For circRNA array experiments (Arraystar, Rockville, MD, USA), AAA tissue specimens were collected from patients who underwent elective surgical repair (n=11), while healthy aortic fragments were harvested from kidney transplant donors and used as controls (n=6). Upon RNA extraction and quality control, samples were treated with RNAseR (Epicentre, Teddington, UK), and reverse transcribed by using fluorescently labeled random primers. The resulting labelled cDNA was then purified and 1ug was fragmented, heated, and subsequently hybridized with an 8*15K commercially available array chip displaying 13,617 human circRNAs (Arraystar, #AS-S-CR-H-V2.0) for 17 hours at 65°C in an Agilent Hybridization Oven. Array probes were designed on the backsplicing junction. After washing of slides, the arrays were scanned by the Agilent Scanner G2505C. Agilent Feature Extraction software (version 11.0.1.1) was used to analyze acquired array images. Quantile normalization and subsequent data processing were performed using the R software limma package.

Significantly differentially expressed circRNAs between two groups were defined as having an absolute fold change > 1.5 and p-value < 0.05. Based on these conditions, gene overexpression analysis of Gene Ontology terms was performed, based on the categorization of their linear RNA counterparts.

### Primary cell culture of human aortic aneurysm VSMCs

Written and informed consent was obtained from all patients, and protocols were approved by the local ethics committee. Human AAA specimens were harvested during surgical repair and stored in complete DMEM/F12 Medium containing 5% FBS and 1% PBS (Millipore, Darmstadt, Germany). The tissue was placed in a sterile Petri dish and washed with PBS. Adventitia, neo-intima, and excessive calcification were removed, and the remaining tissue was cut into small pieces using a sterile scalpel. Enzymatic digestion was carried out in complete DMEM/F12 medium implemented with 1.4 mg/mL Collagenase A (Roche, Penzberg, Germany), for 4–6 hours in a humidified incubator at 37°C and 5% CO2. Cells were strained using a 100μm cell strainer to remove debris. After 2 washings, cells were resuspended in 7 mL complete DMEM/F12 Medium and placed in culture in a small cell culture flask in a humidified incubator at 37°C and 5% C. Medium was changed every other day. After seven days, the medium was replaced with SMC Growth Medium (PeloBiotech, Planegg, Germany). Cells were used between passages 3 and 11. Primary AoSMCs from healthy donors (control AoSMCs, CTRL) were purchased from Cell Applications (#354-05a) and cultured in SMC Growth Medium, following manufacturer’s instructions. Cells were used between passages 5 and 7.

### Validation of circular junctions

For circRNA analysis, RNAseR (Lucigen, USA) was applied to extracted RNA at a concentration of 2U/ug RNA for 15 minutes, followed by enzyme inactivation 20 minutes at 65°C and column-based purification (Qiagen, #74204). First strand cDNA synthesis was performed with the High-Capacity-RNA-to-cDNA Kit (Applied Biosystems, Waltham, MA, USA) according to manufacturer’s instructions, starting from equal amounts of purified RNA. Divergent primers were designed to amplify the circular junction by taking advantage of circInteractome tool (https://circinteractome.nia.nih.gov/). Oligonucleotides sequences and provider details are reported in **Table S2**. PCR products were visualized using a FastGene UV Transilluminator (Nippon Genetics, Düren, Germany) and individual bands were gel extracted (Qiagen, Hilden, Germany) and subsequently cloned on a TOPO TA Cloning Kit (Invitrogen) for Sanger Sequencing (Eurofins Genomics, Ebersberg, Germany).

### Silencing, transfection, and doxorubicin treatment of human aortic smooth muscle cell

CircRNA/ mRNA silencing was performed by using Lipofectamine RNAiMAX (Thermo Fisher Scientific, Waltham, MA) reagent, according to manufacturer’s instructions. siRNAs targeting the backsplicing junction were designed with the help of CircInteractome tool (https://circinteractome.nia.nih.gov/) and used at a final concentration of 100nM for 48 or 72 hours. A scrambled siRNA (Ambion) was employed as negative control. A complete list of siRNAs sequences is reported in **Table S3**. For doxorubicin treatment, 10uM doxorubicin (SIGMA, D1515) or DMSO (SIGMA, D8418) in cell medium was administered to cells for 24h. Phosphorylation of p53 was used as treatment readout.

### RNA isolation and gene expression analysis

Total RNA was isolated with a Qiazol-based (Qiagen, Hilden, Germany) RNA isolation protocol, by taking advantage of miRNeasy Mini (for tissue) or Micro (for cells) Kit (Qiagen, Hilden, Germany). RNA was quantified by NanoDrop (Wilmington, DE, USA) and RNA quality was verified with Agilent 2100 Bioanalyzer (Agilent Technologies, Santa Clara, CA, USA). DNAse (Qiagen, Hilden, Germany) was applied to avoid artifacts deriving from DNA contamination. First strand cDNA synthesis was performed with the High-Capacity-RNA-to-cDNA Kit (Applied Biosystems, Waltham, USA) according to manufacturer’s instructions, starting from equal amounts of purified RNA. Quantitative Real-time PCR (qPCR) was performed on a QuantStudio3 Real-Time PCR System (Applied Biosystems, Waltham, USA), by using Sybr-Green PCR Master Mix (Roche, Planegg, Germany) or TaqMan™ Fast Advanced Master Mix (Applied Biosystems, Waltham, USA). Oligonucleotide sequences/Taqman assays used in this paper are listed in **Table S2**. Amplified transcripts were quantified by using the comparative Ct method and relative gene expression calculated by the method of ΔΔCt ^42^ and are expressed as mean ± SEM. All experiments included at least 3 replicates per group.

### RNA isolation from serum

RNA was isolated from 9 AAA and 10 PAD patients’ serum using the miRNeasy Serum/Plasma Advanced Kit (Qiagen, Hilden, Germany), following in principle the manufacturer’s instructions. Briefly, 1.25μl of MS2 RNA (Roche, Planegg, Germany) and 100ng of IVT *GFP* RNA (as spike-in control) were combined with 500μl of serum. After adding 240μl of Buffer RPL, samples were incubated at room temperature for 3 minutes. 80μl Buffer RPP were further added and incubated for 3 min. After centrifugation, the supernatant was mixed with 1 volume isopropanol and transferred to a RNeasy UCP MinElute column. RWT and RPE washings were eventually followed by RNA precipitation in 80% ethanol. Elution was carried out in 20μl RNase-free water.

### Single-cell RNA Sequencing (scRNA-seq)

Human aneurysmal abdominal arteries were harvested during open repair in the Department of Vascular and Endovascular Surgery at the Klinikum rechts der Isar of the Technical University Munich. After enzymatic digestion, tissue dissociation was performed by using the Multi Tissue Dissociation Kit 2 (Miltenyi Biotech, 130-110-203), GentleMACS Dissociator (Miltenyi Biotech, 130-093-235), GentleMACS C tubes (Miltenyi Biotech, 130-096-334), and the 37C_Mulit_G program, all according to the manufacturer’s instructions. The cell suspension was strained (70 µm, 40 µm) and Dead Cell Removal (Miltenyi Biotech, 130-090-101) using MS Columns (Miltenyi Biotech, 130-042-201) was performed. Cells were resuspended in PBS + 0,04% BSA. Cells were loaded into a 10x Genomics microfluidics Chip G and encapsulated with barcoded oligo-dT-containing gel beads by utilizing the 10x Genomics Chromium Controller. Gel Beads-in-emulsion (GEM) clean-up, cDNA Amplification and 3’Gene Expression Dual Index Library Construction was performed according to the manufacturer’s instructions (CG000315 Rev C). Sequencing was performed by taking advantage of an Illumina NovaSeq 6000 Sequencing system. Data relative to N=4 AAA patients’ specimens was used to perform the downstream analysis. Raw data obtained from sequencing was demultiplexed according to the 10x Genomics pipeline by using Cell Ranger v2.1.0 software (https://support.10xgenomics.com). A gene-barcode matrix was generated for each library, and cell barcodes and UMIs were corrected and filtered. Single-Cell Data Analysis R package Seurat (version 4.1.1)^43,44^ was used for the analysis in RStudio (version 1.4.1717). The CellCycleScoring function^45^ in Seurat was applied to calculate the scores of cell cycle phases. For this, the following filter conditions were applied: nFeature_RNA= more than 4000 and less than 100; nCount= more than 20000 and percent_mt= more than 15%. nFeature_RNA represents the number of all genes detected in each cell; nCount_RNA represents the sum of the expression level of all genes in each cell; and percent.it represents the proportion of mitochondrial genes detected in each cell. A total of 4377 cells were used for the downstream analysis. The transform normalization workflow^46^ was adopted to mitigate possible technically driven or other variations, in which mitochondrial genes and cell cycle phase were regressed out. UMAP (Uniform Manifold Approximation and Projection) was used for converting cells into two-dimensional maps. The FindAllMarkers function was performed to detect the main features of each cluster with default parameters. The top expressed genes were used for cell type identification.

### Bulk sequencing of AAA patient- and CTRL donor-derived AoSMCs

Cells were collected in Qiazol (Qiagen, Hilden, Germany) and RNA isolated by taking advantage of miRNeasy Mini Kit (Qiagen, Hilden, Germany). RNA was quantified by NanoDrop (Wilmington, DE, USA), and quality control was performed with Agilent 2100 Bioanalyzer (Agilent Technologies, Santa Clara, CA, USA). Sequencing libraries preparation was performed using TruSeq stranded total RNA kit (Illumina). Between 35 and 40 million RNA-seq 150 bp pair-end reads were generated for each sample. Quality control (QC) was performed on both raw sequencing files, and alignment performance was generated after mapping read pairs to the mouse reference genome; the quality was consistently high throughout these early data processing stages. The metadata and metrics related to sequencing depth and alignment to the reference genome were then assessed for confounding factors. Following gene expression quantification, lowly expressed genes were excluded, and exploratory data analysis (EDA) was performed on the samples pre- and post-normalization. Differential expression and functional enrichment analysis were defined based on expression fold changes (FC) between the patient-derived and control cell lines at FC ≥ 16 (log2(FC) ≥ 4, indicating very strong differential expression). Gene set enrichment analysis of Reactome pathways and Gene Ontology (GO) terms was eventually performed.

### Double immunofluorescence staining

Human AAA/ control tissue samples were mounted on poly-L-lysine pre-coated SuperFrost Plus slides (Thermo Fisher Scientific). Human OCT-embedded frozen tissue was cut into 8μm thick slides, dried and stored at -80°C. Fixation was performed in ice-cold acetone for 10 minutes. For paraffin-embedded samples, fixation was performed for 48 hours in 4% paraformaldehyde at room temperature and sections of 3μm were cut. Only for these samples, deparaffination and antigen retrieval were performed. After blocking of peroxidase activity (0.3% hydrogen peroxide for 15minutes), additional blocking with 5% horse serum was performed for 1 hour. Primary and secondary antibodies were diluted in 5% horse serum as follows: anti-ATM ab32420 1:100; anti-SMA ab78171:200. Both primary antibodies were incubated after one-another overnight at 4°C, followed by secondary antibodies for 1 hour each at the respective day. For negative controls, only the secondary antibody was applied for 1 hour at room temperature. TrueBlack Lipofuscin Autofluorescence Quencher (Biotium, Fremont, CA, USA) was applied to reduce background fluorescence. Sections were counterstained with DAPI (Thermo Fisher Scientific, Waltham, MA, USA), and images were taken under a confocal microscope (Olympus FV3000).

### Immunohistochemistry (IHC)

2µm sections of paraffin-embedded tissue were mounted on SuperFrost slides (Thermo Fisher Scientific, Waltham, MA, USA) and standard hematoxylin-eosin (HE) and Elastica van Gieson (EVG) stainings were performed. For immunohistochemistry (IHC), sections were mounted on 0.1% poly-L-lysine (Sigma-Aldrich, St. Louis, MO, USA) pre-coated SuperFrost Plus slides (Thermo Fisher Scientific, Waltham, MA, USA). For antigen retrieval, slides were boiled in a pressure cooker with 10nM citrate buffer (distilled water with citric acid monohydrate, pH 6.0), and endogenous peroxidase activity was blocked with 3% hydrogen peroxide. Consecutive slides were incubated with anti-ATM (1:100) or anti-SMA (1:200) diluted in Dako REAL Antibody Diluent (Dako, Glostrup, Denmark). Slides were then treated with biotinylated secondary antibodies and target staining was performed with peroxidase-conjugated streptavidin and DAB chromogen (Dako REAL Detection System Peroxidase/DAB+, Rabbit/Mouse Kit; Dako, Glostrup, Denmark). Mayer’s hematoxylin (Carl Roth, Karlsruhe, Germany) was used for counterstaining and appropriate positive and negative controls were performed for each antibody. All slides were scanned with an Aperio AT2 (Leica, Wetzlar, Germany), and images were taken with the Aperio ImageScope software (Leica, Wetzlar, Germany).

### *In situ* hybridization (ISH)

Qiagen miRCURY locked nucleic acid DIG (digoxigenin)-labeled probes (sense c*ATM*-DIG: 5’DIG-AGTGGTTAGACAGTGATGTGT-DIG 3’) (Qiagen, Hilden, Germany) were used for ISH, performed according to manufacturer’s instructions. A complete list of ISH probes sequences is reported in **Table S2**. Briefly, tissue sections were either de-paraffinized (formalin-fixed paraffin-embedded) or thawed (frozen) and rehydrated. After proteinase K permeabilization, cATM probe hybridization was carried out at 54°C for 2 hours in a hybridization oven. A negative control (no probe) was performed in parallel. Slides were washed in pre-wormed saline-sodium citrate buffers at hybridization temperature, with subsequent DIG detection methods as previously described^47^. Nuclear counterstaining was performed with Nuclear Fast Red (Sigma-Aldrich, Saint Louis, MO, USA).

### Protein isolation and Western blotting

Cells were homogenized in RIPA buffer (Thermo Fisher Scientific, Waltham, MA, USA) including protease and phosphatase inhibitor cocktails (Sigma-Aldrich, Saint Louis, MO, USA). Protein concentrations were determined by using the Bicinchoninic Acid assay (Thermo Fisher Scientific, Waltham, MA, USA), following the manufacturer’s protocol. Protein samples (10-40μg/well) were mixed with LDS 4X Sample Buffer (Novex) and Sample reducing agent (Novex), denatured at 95°C for 5 minutes and loaded on NuPage™ 3-8% gels. Following electrophoresis and electrotransfer, blots were blocked with 5% milk in Tris-buffered saline + 0,1% Tween-20 and probed with specific antibodies diluted in blocking solution. Signals were revealed after incubation with horseradish peroxidase (HRP)-conjugated secondary antibodies (Abcam) 1:10000, in combination with ECL (GE Healthcare). Image detection was performed with C600 Azure Biosystems Imager (Biozym)/ChemiDoc XRS System (Bio-rad, Hercules, CA, USA) and image quantification was carried out with ImageJ software.

Employed antibodies were all diluted in 5% milk in TBS-T as follows: ß-actin #A1978 (Sigma, USA), 1:10000; ATM **#**ab3242 1:500; p53 #ab131442 1:1000; phospho-p53 #ab1431 1:500; phospho-ATM #ab81292 1:500; MY11C #ab53219 1:500; ß-Tubulin #ab6046; Caspase 3 #ab13837, 1:250, Active Caspase 3 #5A1E, Cell Signalling).

### Cell fractionation

Nucleocytoplasmic fractionation was performed as previously described in^48^. Fractions were extracted from confluent hAoSMCs cultured in T75cm2-flasks (Corning) and RNA isolated using Qiazol reagent, as explained above. The purity of the nuclear and cytoplasmic fractions was confirmed by real-time quantitative PCR on *GAPDH/RPLPO/ß Actin* and *NEAT1*, respectively.

### Kinetic assessment of proliferation and apoptosis in human AoSMCs

Real-time assessment of hAoSMCs status was carried out with an IncuCyte Zoom System (Sartorius, Goettingen, Germany), as previously described by our group ^49,50^. Live cell imaging was performed upon target silencing or doxorubicin treatment. To evaluate the death rate, a Caspase 3/7 Apoptosis reagent (Sartorius, Goettingen, Germany) was added at a final concentration of 5μM prior to imaging with phase contrast/ fluorescence (4h/imaging pattern). Images were auto-collected and analyzed by using the IncuCyte software package.

### ATM enzyme-linked immunoassay (ELISA)

ATM ELISA was performed on 7 PAD *vs*. 7 AAA patient’s serum samples, according to manufacturer’s instructions (NBP2_69891, Novus Biologicas, USA). Briefly, 80μl of standard or serum were aliquoted into anti-ATM-coated wells in duplicate and incubated. After application of biotinylated antibody, wells were washed and provided with HRP conjugate. After further washings, the signal was revealed *via* substrate addition and stopped after 15 minutes. Absorbance was read at 450nm with a plate reader (Molecular Devices, Germany).

### Statistical analysis

All data are expressed as means ± sd for n ≥ 3 replicas. Statistical analysis was performed using Graphpad Prism software. Statistically significant differences were assessed by unpaired Student’s t test/ Mann-Whitney-U-Test. Values of P < 0.05 were considered significant.

## DATA AVAILABILITY

Circular RNA array and scRNA-Seq data can be provided by the corresponding author upon reasonable request.

## ACKNOWLEDGMENTS

We are indebted to all the members of the Molecular Vascular Medicine lab at the Technical University Munich for thought-provoking discussions. We thank Renate Hegenloh and the Munich Vascular Biobank team for technical support with human specimens. This work was supported by funding from the Swedish Heart-Lung-Foundation (20210450), the Swedish Research Council (Vetenkapsrådet, 2019-01577), a DZHK Translational Research Project on microRNA modulation in aortic aneurysms, the CRC1123 and TRR267 of the German Research Council (DFG), the National Institutes of Health (NIH; 1R011HL150359-01), and the Bavarian State Ministry of Health and Care through the research project DigiMed Bayern.

## AUTHOR CONTRIBUTIONS

L.M. and F.F. conceived and designed the project and wrote the manuscript. F.F. supervised the research, executed the experiments, and analysed and interpreted the data. G.W. designed and executed the experiments, curated, and analysed the data, and participated in data interpretation. Z.W., Z.L., H.W, J.P., R.B., V.P., and N.G assisted with the experiments and data collection and analysed the data. H.H.E., A.B., C.K. and N.S. collected and provided human specimens (Munich Vascular Biobank). R.H. and J.R. collected and provided human specimens (Stockholm Aortic Aneurysm Biobank).

---

**Table S1.** Gene set enrichment analysis. The top 30 significantly enriched Gene Ontology (GO) terms and Reactome pathways terms in AAA patient-derived over CTRL AoSMCs are shown. Selected pathways and functions are ranked by FDR-adjusted p-value (qvalue). For each pathway and GO function tested, the following statistics were retrieved: GeneRatio, pvalue, qvalue (FDR-adjusted p-value) and Count. ‘Count’ corresponds to the number of pathway/function-specific genes that were assessed. ‘GeneRatio’ represents the number of genes enriched in this pathway among all differentially expressed genes.

| Pathway | GeneRatio | pvalue | qvalue | Count |
| --- | --- | --- | --- | --- |
| Translation | 287/8140 | 4.73696E-31 | 3.71976E-28 | 287 |
| Transcriptional Regulation by TP53 | 349/8140 | 5.06615E-27 | 1.98913E-24 | 349 |
| Organelle biogenesis and maintenance | 282/8140 | 3.24234E-21 | 8.48695E-19 | 282 |
| Processing of Capped Intron-Containing Pre-mRNA | 236/8140 | 7.23595E-21 | 1.42053E-18 | 236 |
| mRNA Splicing | 188/8140 | 3.54202E-20 | 5.56284E-18 | 188 |
| RHO GTPase cycle | 405/8140 | 1.51346E-19 | 1.98077E-17 | 405 |
| mRNA Splicing - Major Pathway | 180/8140 | 3.26217E-19 | 3.65952E-17 | 180 |
| rRNA processing | 198/8140 | 2.63257E-18 | 2.58408E-16 | 198 |
| rRNA processing in the nucleus and cytosol | 189/8140 | 3.35583E-18 | 2.92801E-16 | 189 |
| Major pathway of rRNA processing in the nucleolus and cytosol | 179/8140 | 4.84223E-17 | 3.80242E-15 | 179 |
| S Phase | 159/8140 | 1.06731E-16 | 7.61924E-15 | 159 |
| Regulation of TP53 Activity | 157/8140 | 1.84689E-16 | 1.20858E-14 | 157 |
| Mitotic Metaphase and Anaphase | 224/8140 | 8.86736E-16 | 5.35632E-14 | 224 |
| Mitotic Anaphase | 223/8140 | 1.12525E-15 | 6.31155E-14 | 223 |
| DNA Repair | 306/8140 | 6.85223E-15 | 3.5872E-13 | 306 |
| Influenza Infection | 152/8140 | 7.32073E-15 | 3.59294E-13 | 152 |
| Influenza Viral RNA Transcription and Replication | 133/8140 | 1.07966E-14 | 4.98716E-13 | 133 |
| tRNA processing | 107/8140 | 3.49832E-14 | 1.52617E-12 | 107 |
| Mitotic G1 phase and G1/S transition | 145/8140 | 4.7199E-14 | 1.80204E-12 | 145 |
| Mitotic G2-G2/M phases | 188/8140 | 4.84368E-14 | 1.80204E-12 | 188 |
| Cell Cycle Checkpoints | 270/8140 | 4.87095E-14 | 1.80204E-12 | 270 |
| HIV Infection | 219/8140 | 5.04862E-14 | 1.80204E-12 | 219 |
| Mitotic Prometaphase | 193/8140 | 6.89578E-14 | 2.35435E-12 | 193 |
| Diseases of signal transduction by growth factor receptors and second messengers | 354/8140 | 7.51988E-14 | 2.46045E-12 | 354 |
| G2/M Transition | 186/8140 | 7.85667E-14 | 2.46782E-12 | 186 |
| Asparagine N-linked glycosylation | 278/8140 | 8.4857E-14 | 2.56289E-12 | 278 |
| Antigen processing; Ubiquitination & Proteasome degradation | 281/8140 | 3.84092E-13 | 1.11238E-11 | 281 |
| Signaling by ROBO receptors | 204/8140 | 3.96641E-13 | 1.11238E-11 | 204 |
| G1/S Transition | 128/8140 | 4.87919E-13 | 1.32119E-11 | 128 |
| Cellular response to starvation | 150/8140 | 6.73634E-13 | 1.7239E-11 | 150 |

| **GO terms** | **GeneRatio** | **pvalue** | **qvalue** | **Count** |
| --- | --- | --- | --- | --- |
| focal adhesion | 400/13332 | 3.79307E-47 | 1.27766E-44 | 400 |
| cell-substrate junction | 404/13332 | 4.12366E-45 | 6.94512E-43 | 404 |
| proteasomal protein catabolic process | 455/12835 | 3.72932E-46 | 1.4458E-42 | 455 |
| ncRNA metabolic process | 460/12835 | 5.39947E-44 | 1.04665E-40 | 460 |
| mitochondrial matrix | 444/13332 | 4.43355E-41 | 4.97802E-39 | 444 |
| proteasome-mediated ubiquitin-dependent protein catabolic process | 401/12835 | 2.82198E-41 | 3.64678E-38 | 401 |
| mRNA catabolic process | 358/12835 | 1.76697E-40 | 1.71257E-37 | 358 |
| RNA catabolic process | 390/12835 | 1.12803E-39 | 8.74641E-37 | 390 |
| ncRNA processing | 378/12835 | 1.96832E-39 | 1.27181E-36 | 378 |
| ribosome biogenesis | 298/12835 | 1.24209E-38 | 6.87912E-36 | 298 |
| protein polyubiquitination | 320/12835 | 8.43636E-35 | 4.08831E-32 | 320 |
| ribonucleoprotein complex biogenesis | 432/12835 | 1.85606E-34 | 7.99518E-32 | 432 |
| macroautophagy | 296/12835 | 7.45544E-33 | 2.89035E-30 | 296 |
| covalent chromatin modification | 419/12835 | 3.39542E-32 | 1.19668E-29 | 419 |
| nuclear transport | 322/12835 | 4.6049E-32 | 1.45533E-29 | 322 |
| nuclear-transcribed mRNA catabolic process | 206/12835 | 4.88008E-32 | 1.45533E-29 | 206 |
| histone modification | 408/12835 | 7.1128E-32 | 1.96966E-29 | 408 |
| protein targeting | 402/12835 | 1.18821E-31 | 2.99473E-29 | 402 |
| nucleocytoplasmic transport | 319/12835 | 1.23595E-31 | 2.99473E-29 | 319 |
| cell cycle G2/M phase transition | 265/12835 | 1.33706E-31 | 3.04917E-29 | 265 |
| ubiquitin ligase complex | 276/13332 | 4.13223E-31 | 3.47977E-29 | 276 |
| nuclear speck | 377/13332 | 1.02294E-30 | 6.89137E-29 | 377 |
| mitochondrial inner membrane | 443/13332 | 1.85298E-30 | 1.04027E-28 | 443 |
| Golgi vesicle transport | 344/12835 | 1.94251E-30 | 4.18378E-28 | 344 |
| spindle | 351/13332 | 1.15875E-29 | 5.57593E-28 | 351 |
| rRNA processing | 222/12835 | 9.10313E-30 | 1.85744E-27 | 222 |
| mitotic nuclear division | 280/12835 | 1.05761E-29 | 2.05009E-27 | 280 |
| rRNA metabolic process | 230/12835 | 5.28977E-29 | 9.76552E-27 | 230 |
| G2/M transition of mitotic cell cycle | 247/12835 | 7.18062E-29 | 1.26537E-26 | 247 |
| regulation of response to DNA damage stimulus | 215/12835 | 1.15866E-28 | 1.95302E-26 | 215 |

**Table S2.**
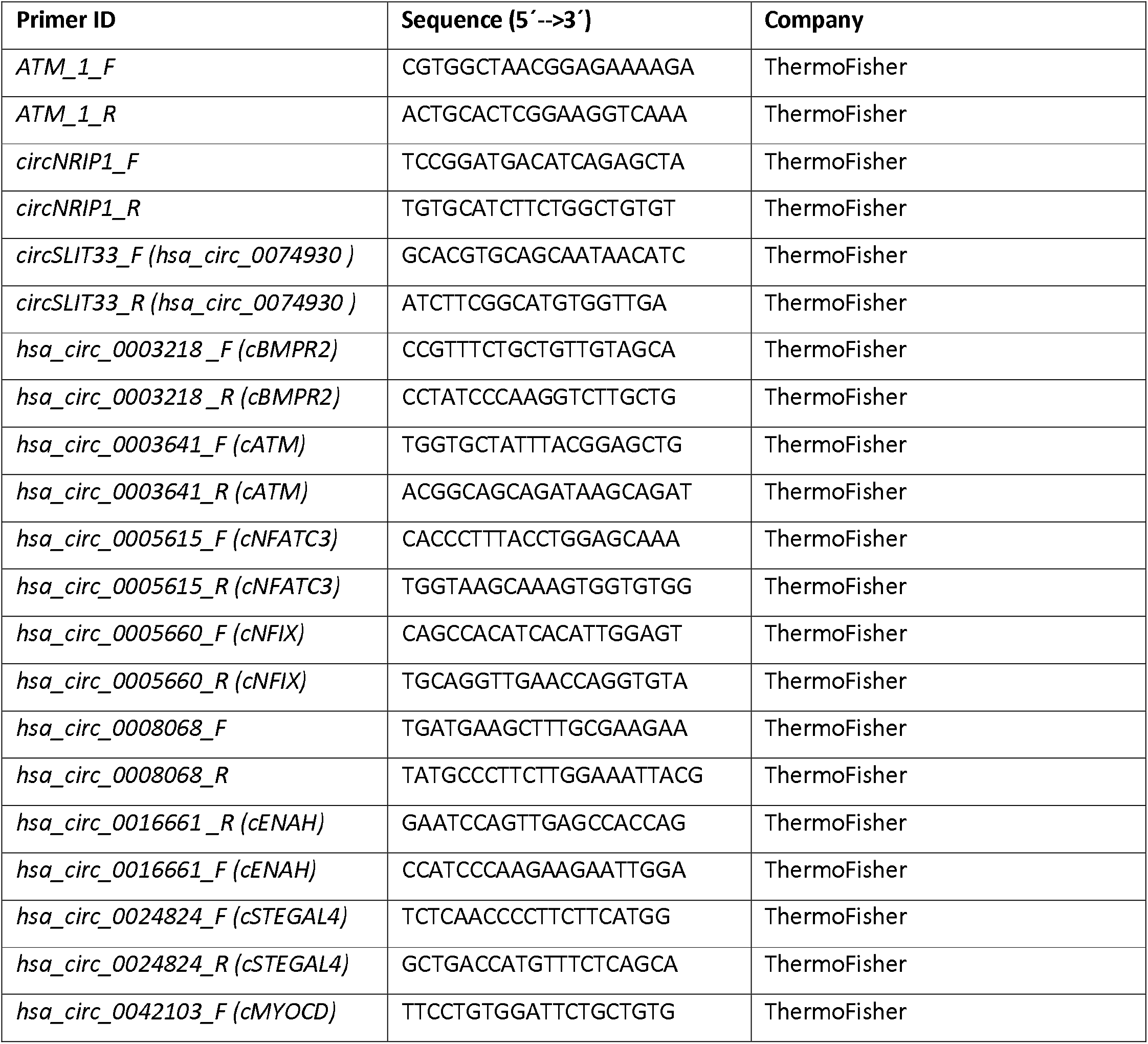
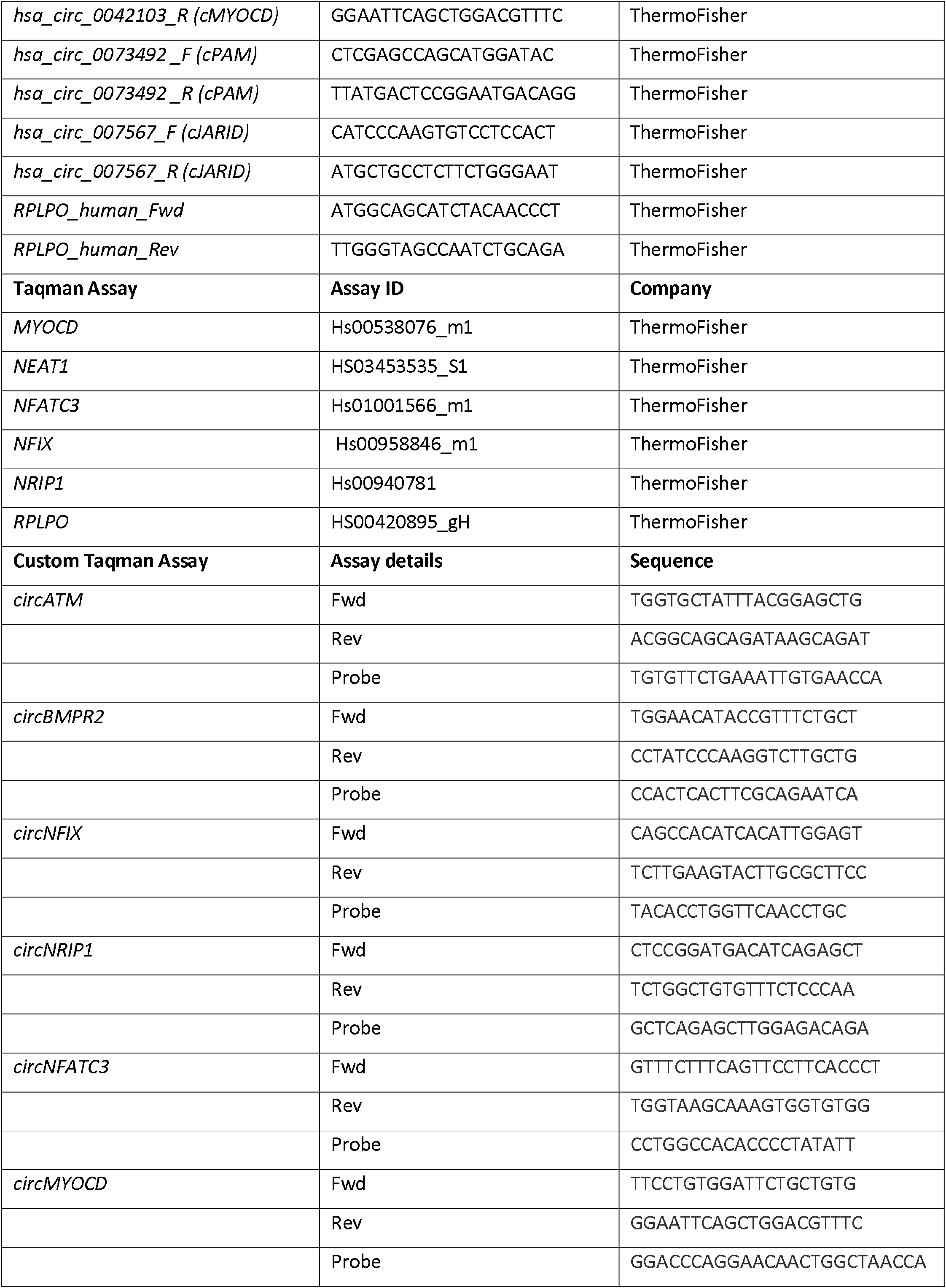
List of PCR oligonucleotides and Taqman assays.

**Table S3.**
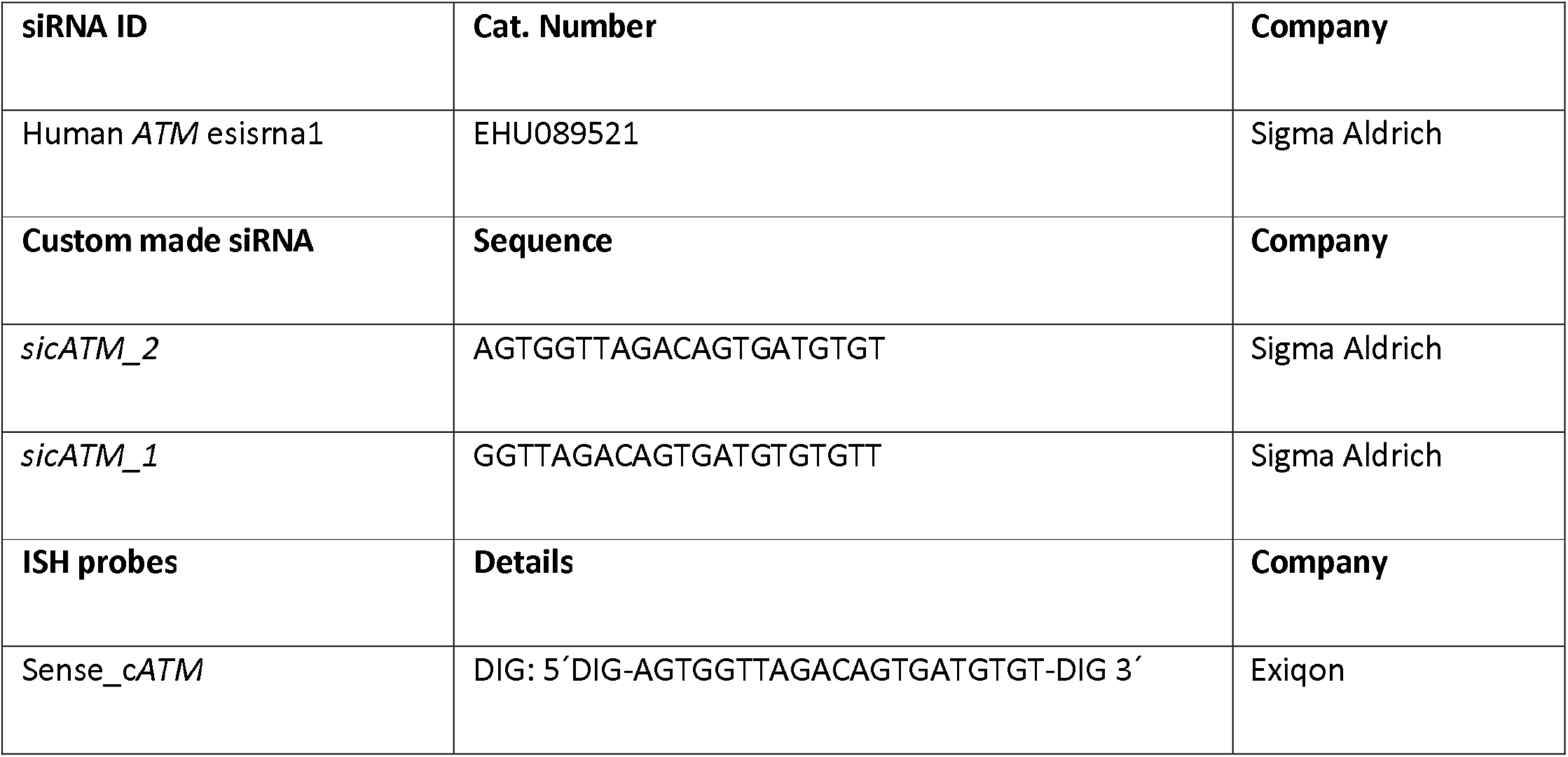
List of oligonucleotides for silencing and *in situ* hybridization.

