## Supplementary Files for "The circular RNA Ataxia-telangiectasia mutated (*cATM*) regulates oxidative stress in smooth muscle cells in expanding abdominal aortic aneurysms"

**A**

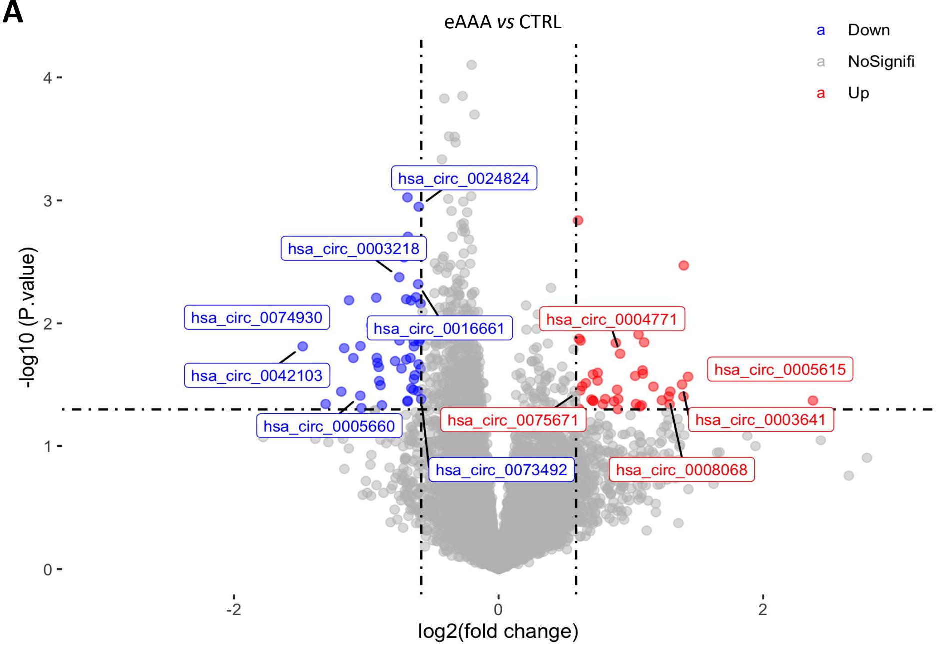


**B**

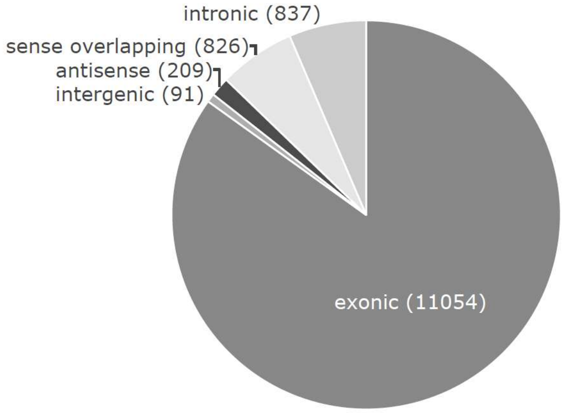


**Figure S1. *Differentially expressed circRNAs in eAAA vs CTRL patients*. (A)** Volcano plot showing up- (red, right) and down- (blue, left) regulated circRNAs in eAAA vs CTRL patients. Vertical dash lines are drawn in correspondence of fold change= 1.5; the horizontal dash line indicates p-value of 0.05. All circRNAs initially meant for validation through PCR with divergent primers are labelled. **(B)** Pie chart illustrating the proportion of exonic, intronic, sense-overlapping and antisense circRNAs in the array. Absolute numbers are further indicated for each group. Abbr.: eAAA: elective AAA; CTRL: control.

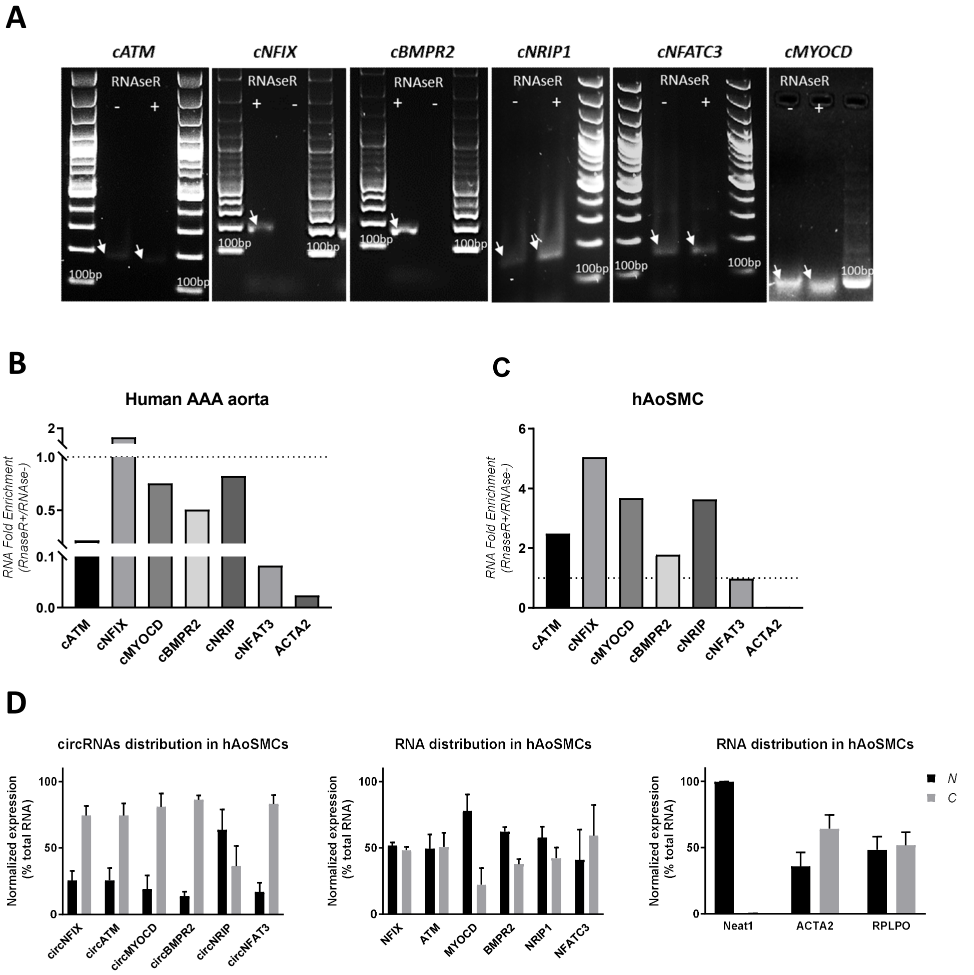


**Figure S2.** ***Validation of circular junctions in human abdominal aortic aneurysm (AAA) tissue specimens and human aortic smooth muscle cells (hAoSMCs).*** **(A)** Agarose gel electrophoresis of circRNAs PCR amplicons showing enrichment upon RNAseR treatment. PCR products were cloned and submitted to Sanger sequencing. RNA fold enrichment upon RNAseR treatment in AAA tissue **(B)** and hAoSMCs **(C)**. Fold enrichment was calculated by comparing CT values in RNAseR+ vs RNAseR- and expressed as 2^-delta CT*100, with delta CT= CT RNAseR+ - CT RNAseR-. Targets displaying values >1 are considered as enriched. **(D)** Subcellular localization of circRNAs (left) and respective linear counterpart (middle) in AoSMCs as quantified by qRT-PCR. Nuclear (abbr.: N) and cytoplasm (abbr.: C) purity was monitored by measuring *ACTA2/ RPLPO* or *NEAT1*, respectively (right). Expression levels are indicated as percentage of total RNA. Data are represented as mean ± SD.

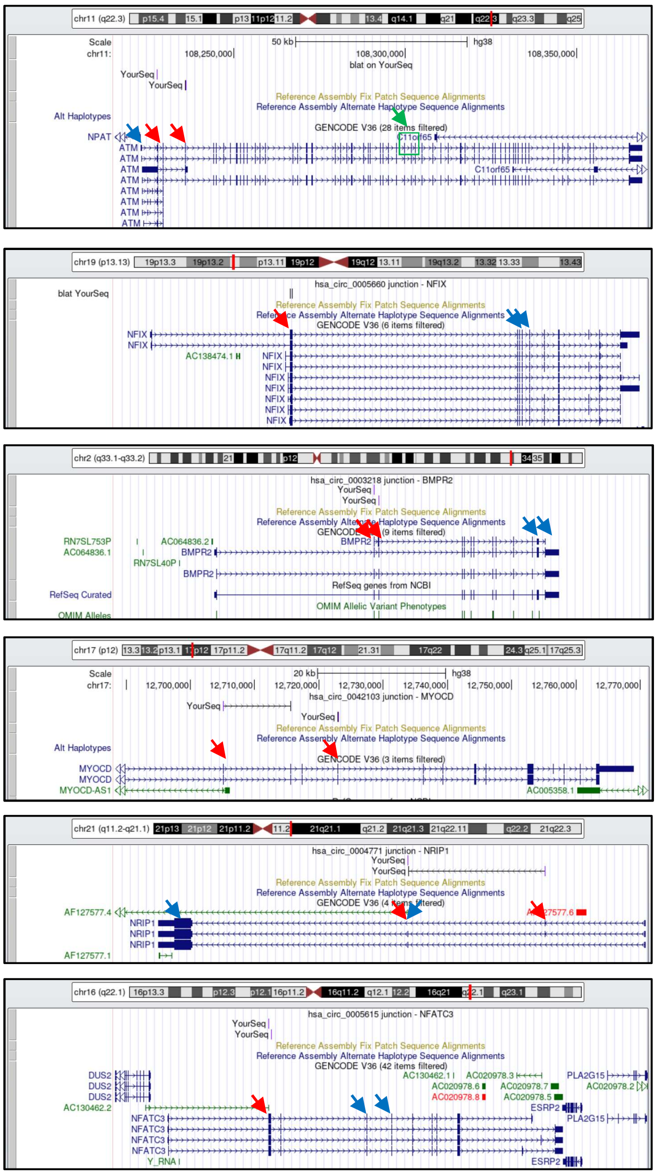

*ATM* **(hsa_circ_0003641)**

*NFIX* **(hsa_circ_0005660)**

*BMPR2* **(hsa_circ_0003218)**

*MYOCD* **(hsa_circ_0042103)**

*NRIP1* **(hsa_circ_0005660)**

*NFATC3* **(hsa_circ_0004771)**

**Figure S3.** ***Genome Browser view of gene loci of validated circRNA targets.*** Red arrows indicate exons involved in backsplicing. Taqman assays and siRNAs targeting circRNAs were design on the backsplicing junction. Taqman assays/ primers for detection of linear transcripts map on blue arrows. SiRNAs target sites of linear transcripts are indicated by green arrows.

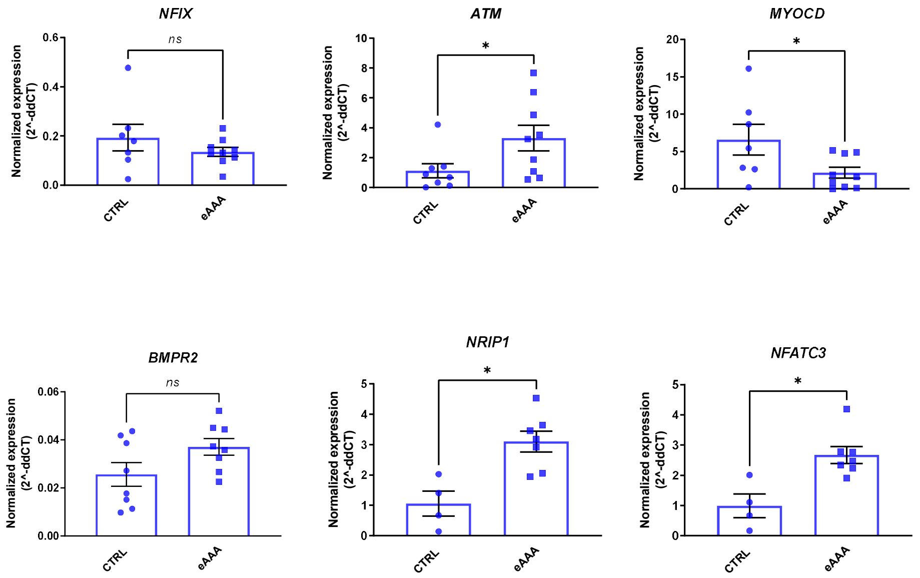


**Figure S4.** ***Expression of linear counterparts of de-regulated circRNAs in eAAA vs CTRL patients.*** **(A)** *ATM, NFIX, MYOCD, BMPR2, NRIP1* and *NFATC3* mRNA levels were determined by qRT-PCR and compared in AAA vs CTRL patients. 2^(-ddCT) was plotted. Data are represented as mean ± SEM. Statistics: T-test. P values < 0.05 were considered as significant. Abbr.: CTRL=control; eAAA: elective AAA.

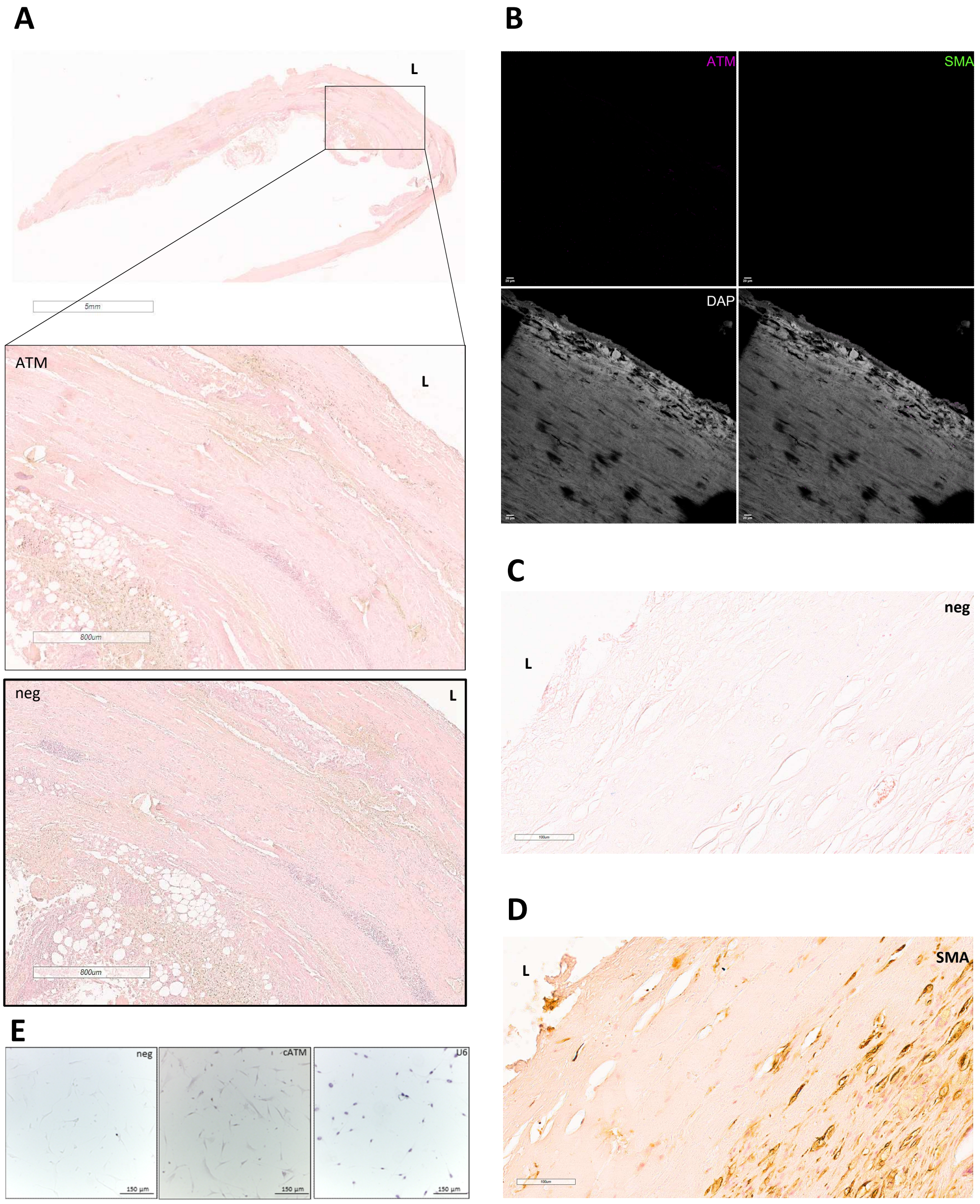


**Figure S5.** ***ATM protein and cATM stainings in human eAAA specimens and aortic SMCs.*** **A.** ATM immunohistochemistry in AAA patient section (top) and relative negative control. **(B)** ATM and SMA IF negative control. **C.** *cATM* ISH negative control and **(D)** SMA IHC staining were performed in consecutive slides. **E.** ISH in human aortic SMCs negative control (left), *cATM* (middle) and U6 positive control (right) signal. L= lumen; neg= negative control

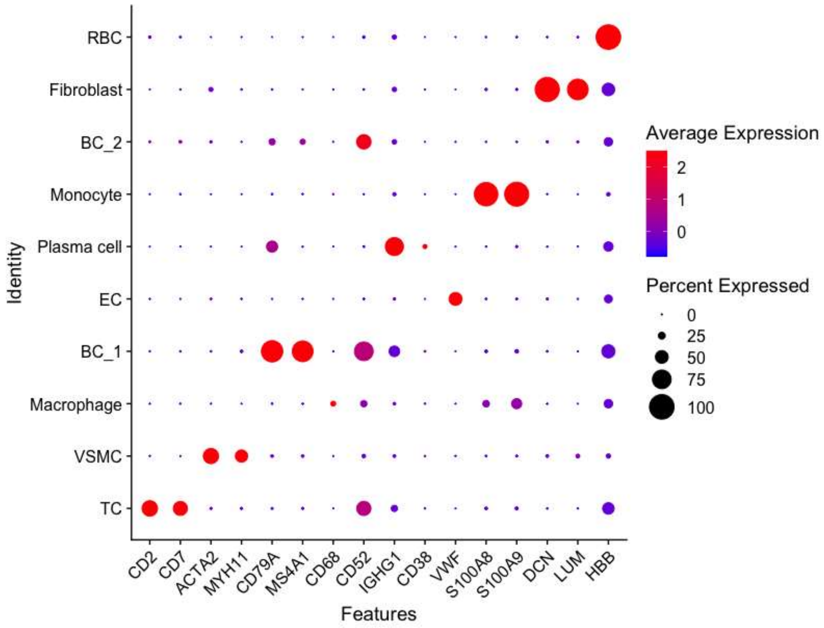


**Figure S6.** ***Clusters identified by single-cell RNA sequencing (scRNA-seq) os human AAA specimens.*** Dot plot depicting the main gene markers employed for cell clusters labeling. Dots colour represents the average expression level (blue=low, red= high) and dots size depicts the percentage of cells expressing the gene in a given cluster.

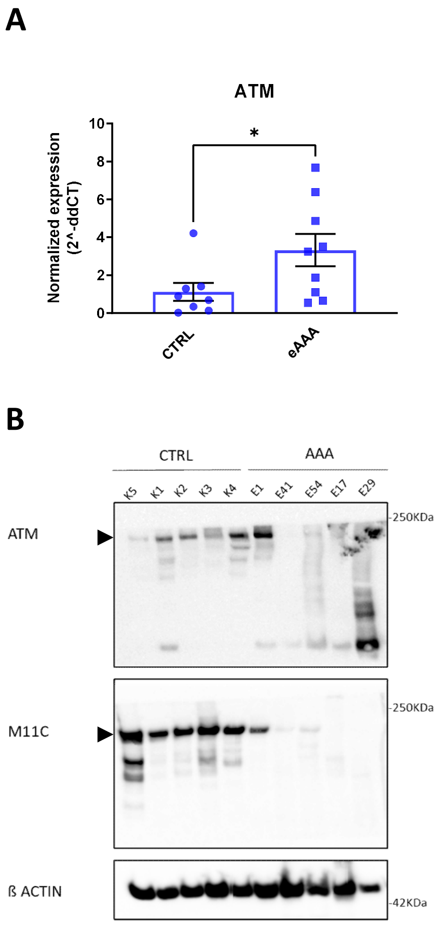


**Figure S7.** ***ATM mRNA and protein expression in eAAA vs CTRL patients*****. A.** *ATM* mRNA levels were determined by qRT-PCR and compared in AAA *vs* CTRL patients. 2^(-ddCT) was plotted. Data are represented as mean ± SEM. Statistics: T-test. P values < 0.05 were considered as significant. **B.** WB showing ATM protein in AAA vs CTRL patients. Black arrows indicate expected molecular weights. Abbr.: L= lumen; neg= negative control CTRL= control; eAAA: elective AAA.

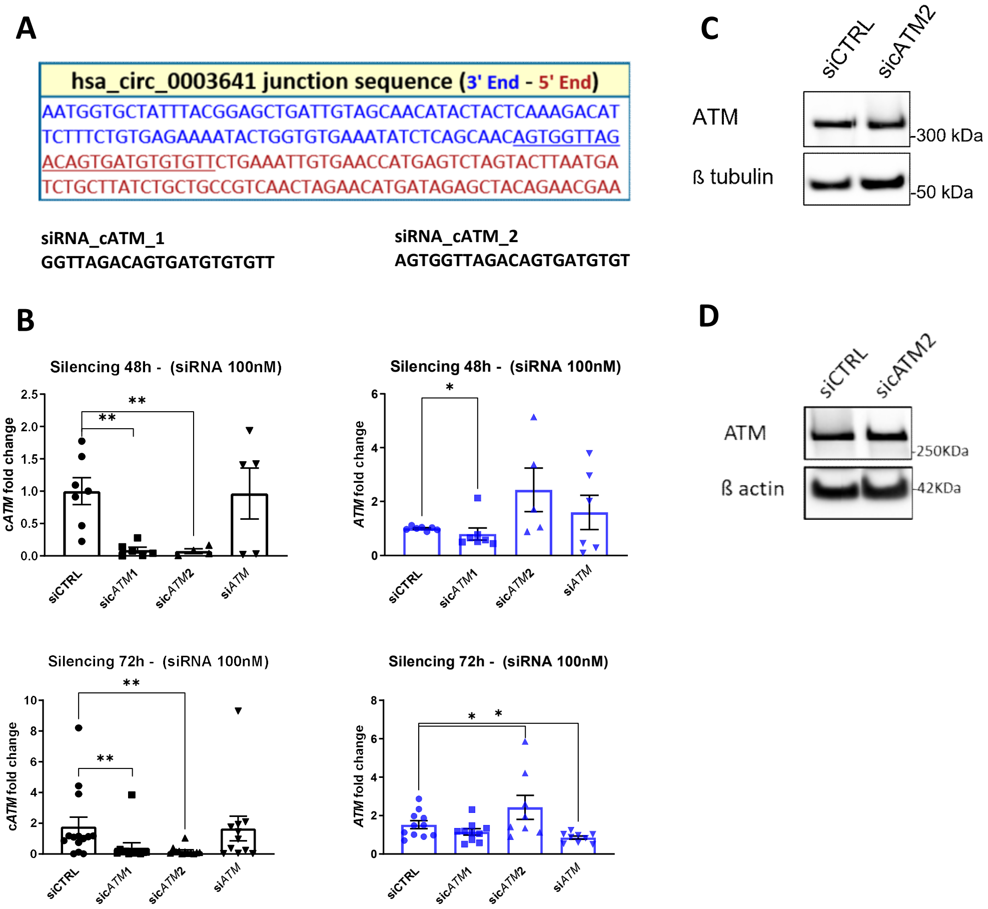


**Figure S8.** ***cATM KD in human aortic smooth muscle cells (hAoSMCs).*** **A.** siRNA design for *cATM* (hsa_circ_0003641). The underlined sequence indicates the region covered by two alternative siRNA (sic*ATM*1 and sic*ATM*2), both centred on the backsplicing junction. Different colours indicate different exons. **B.** qRT-PCR upon c*ATM* and linear *ATM* silencing in control hAoSMCs at 48h (top) and 72h (bottom) with 100nM sic*ATM*1, sic*ATM*2 and si*ATM* and ATM Western Blot **(C)** upon sic*ATM*2 at 72h**.** **D.** ATM Western Blot upon sic*ATM*2 at 72h AAA-derived hAoSMCs.

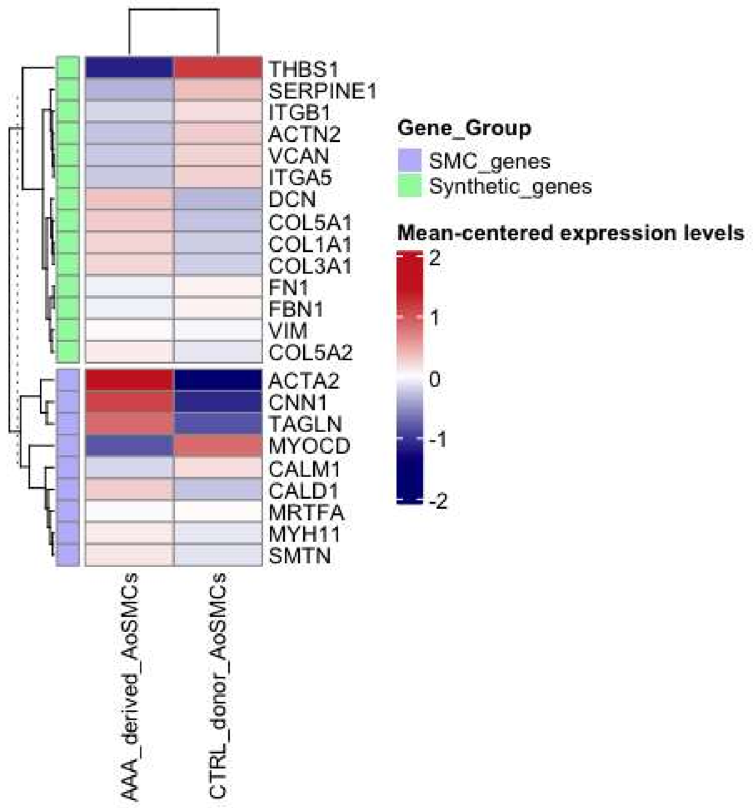


**Figure S9. Gene expression profile of AAA patient-derived *vs* CTRL donor AoSMCs (SMC-characterizing genes). (A)** Heatmap depicts SMC gene signature in AAA *vs* CTRL AoSMCs, as revealed from bulk RNA sequencing (NovaSeq6000). Colours represent the average gene expression level (blue=low, red= high).
